# Highly Constrained Kinetic Models for Single-Cell Gene Expression Analysis

**DOI:** 10.64898/2026.05.22.727214

**Authors:** Hyeon Jin Cho, Christopher H. Bohrer, Pawel Trzaskoma, Jee Min Kim, Aleksandra Pękowska, Rafael C. Casellas, Rob Patro, Carson C. Chow, Daniel R. Larson

## Abstract

Advances in single-cell RNA sequencing (scRNA-seq) and high-resolution imaging techniques, such as single-molecule tracking (SMT) of RNA and transcription factors, allow researchers to quantitatively explore dynamics and variation but have never been integrated into a single coherent model. In this study, we propose a kinetic model that intakes multiple data types, including steady-state and time-resolved datasets, to simulate and fit stochastic models of gene transcription to experimental data. We find that 3-state models provide an essential improvement over the widely used 2-state model for most genes and have the property of kinetic proofreading, which we argue is advantageous in the cellular context. We further identify two dimensionless quantities derived from the rate equations which are broadly conserved across genes. Finally, we extend this model to scRNA-seq datasets to infer kinetic rates under defined perturbations and reveal biochemical insight into the mechanism of action of transcription factors.

## Introduction

Gene expression varies substantially from cell to cell, even among genetically identical cells in a uniform environment. This heterogeneity reflects both structured sources of variation—such as differences in cell state, cell cycle stage, or environmental stimuli—and random fluctuations inherent to biochemical processes [1–5]. Such random variation, or transcriptional noise, arises from the stochastic nature of gene regulation, including the binding of transcription factors (TFs), chromatin remodeling, and mRNA synthesis [1, 6–9]. Consequently, transcription occurs in stochastic bursts, where genes are transcribed in intermittent manner, alternating between active (ON) and inactive (OFF) states [9, 10]. Traditional experimental techniques often mask cellular variability by analyzing the cell population in a bulk condition, assuming uniformity among cells [11]. However, techniques like live-cell imaging combined with fluorescently labeled reporters, single-molecule fluorescent *in situ* hybridization (smFISH), and single-cell RNA sequencing (scRNA-seq) have enabled single-cell measurements of transcriptional activity and mRNA abundance from which transcriptional bursting dynamics can be quantified or inferred [10, 12–14].

Mathematical models of transcriptional dynamics aim to extract interpretable kinetic parameters from observed variability. Among the most widely used is the 2-state model (classic telegraph model) [15], which assumes a gene switches between ON and OFF states and produces mRNA in bursts during the ON state. This model has been further modified and developed into different versions which aim to capture additional biochemical realities and often better represent transcription dynamics [16–21]. However, for some genes, the 2-state telegraph model can oversimplify the dynamics of transcription regulation by excluding intermediate steps that may be partially rate-limiting [22, 23]. Despite its simplicity, the 2-state model has been remarkably effective in describing gene expression across diverse systems, even revealing conserved dynamic control properties [24] and allowing for inference of biochemical rates [25, 26]. Nevertheless, this framework cannot explicitly resolve complex *intermediate* processes between the active and inactive states, such as chromatin remodeling, proximal promoter and enhancer activities and recruitment of general transcription factors (GTFs). In addition, it often fails to capture long and heterogeneous inactive periods observed in experimental data [19, 21]. More complex extensions include 3-state models and kinetic proofreading (KP) models, which introduce additional regulatory checkpoints to capture biophysically plausible steps such as TF and GTF binding or RNA polymerase II (RNAPII) pausing [19, 27, 28]. By fitting these models to single-cell data, researchers aim to distinguish regulatory changes from intrinsic noise, enabling more precise interpretation of perturbation experiments and high-throughput screens such as Perturb-seq [29, 30]. Yet despite increasing use, few modeling frameworks have been rigorously validated by cross-referencing kinetic parameters inferred from smFISH or scRNA-seq with those from live-cell imaging under matched experimental conditions [18, 30, 31].

In this study, we develop and validate a kinetic modeling framework that integrates steady-state transcript distributions from smFISH or scRNA-seq with the dynamic information from live-cell nascent mRNA imaging and single-molecule tracking (SMT). We focus on transcriptional bursting and the role of kinetic proofreading to estimate rates of gene activation, elongation, synthesis, and splicing. We use MCMC sampling to infer gene-specific kinetic rates and perform Bayesian model comparison [32]. To ensure biological fidelity, we constrain our models using experimental measurements of GTF dwell times measured in living cells. This approach allows us to cross-validate parameter estimates across modalities and reveals that the kinetic proofreading model provides a superior fit for a subset of genes imaged live and profiled by smFISH. We then extend this framework to high-throughput scRNA-seq datasets, including the recently published TF Atlas [33], and identify TFs, GTFs, and chromatin remodelers whose overexpression alters kinetic parameters in a manner consistent with proofreading modulation. Overall, our analysis suggests that 3-state models with kinetic proofreading properties are the preferred models of transcriptional regulation, and this experimental approach offers a principled path forward for interpreting transcriptional dynamics in large-scale single-cell datasets.

## Results

### Multimodal gene expression measurements constrain dynamic models

Overall, our theoretical approach is built upon multi-modal experimental measurements of single-cell gene expression carried out in human bronchial epithelial cells (HBEC). This non-transformed cell line is immortalized through CDK4 and hTert over-expression [34] and is tractable for both live-cell imaging and gene editing [21, 35]. These experiments range from low-throughput, high-temporal-resolution characterization of genes by live-cell RNA imaging, to high-spatial-resolution single-molecule snapshot measurements by smFISH, and ultimately to high throughput, transcriptome-wide scRNA-seq.

To capture the temporal dynamics of transcription, we incorporated two types of live-cell imaging datasets: (1) fluorescent tagging of nascent mRNA to track transcriptional bursting dynamics (Figure 1A) and (2) SMT of GTF dwell times (Figure 1B). Temporal data are critical for accurate kinetic modeling of gene regulation, as transcription is inherently a time-dependent and dynamic, multistep process involving GTF binding, transcriptional machinery assembly, promoter activation, RNAPII recruitment and pause release, elongation, splicing, and RNA degradation [5, 10, 36]. Live-cell imaging of nascent mRNA allows quantification of transcription-site fluorescence over time, from which ON/OFF bursting states can be inferred, whereas SMT of GTFs reveals their residence time, providing mechanistic insight into binding specificity and search kinetics [37]. The live-cell nascent RNA imaging dataset contained 10 monoclonal cell lines with MS2 stem loops integrated into introns at endogenous locations, as described previously [21]. These genes are: *CANX, DNAJC5, ERRFI1, KPNB1, MYH9, RAB7A, RHOA, RPAP3, SEC16A,* and *SLC2A1*. For the GTF SMT imaging dataset, we analyzed the binding and unbinding dynamics of TATA box binding protein (TBP), a core component of TFIID, as described previously [38]. TBP is thought to have functional interactions with thousands of genes [39, 40], making it a good candidate for a generalized model of transcription, as discussed in more detail below.

**Figure 1.**
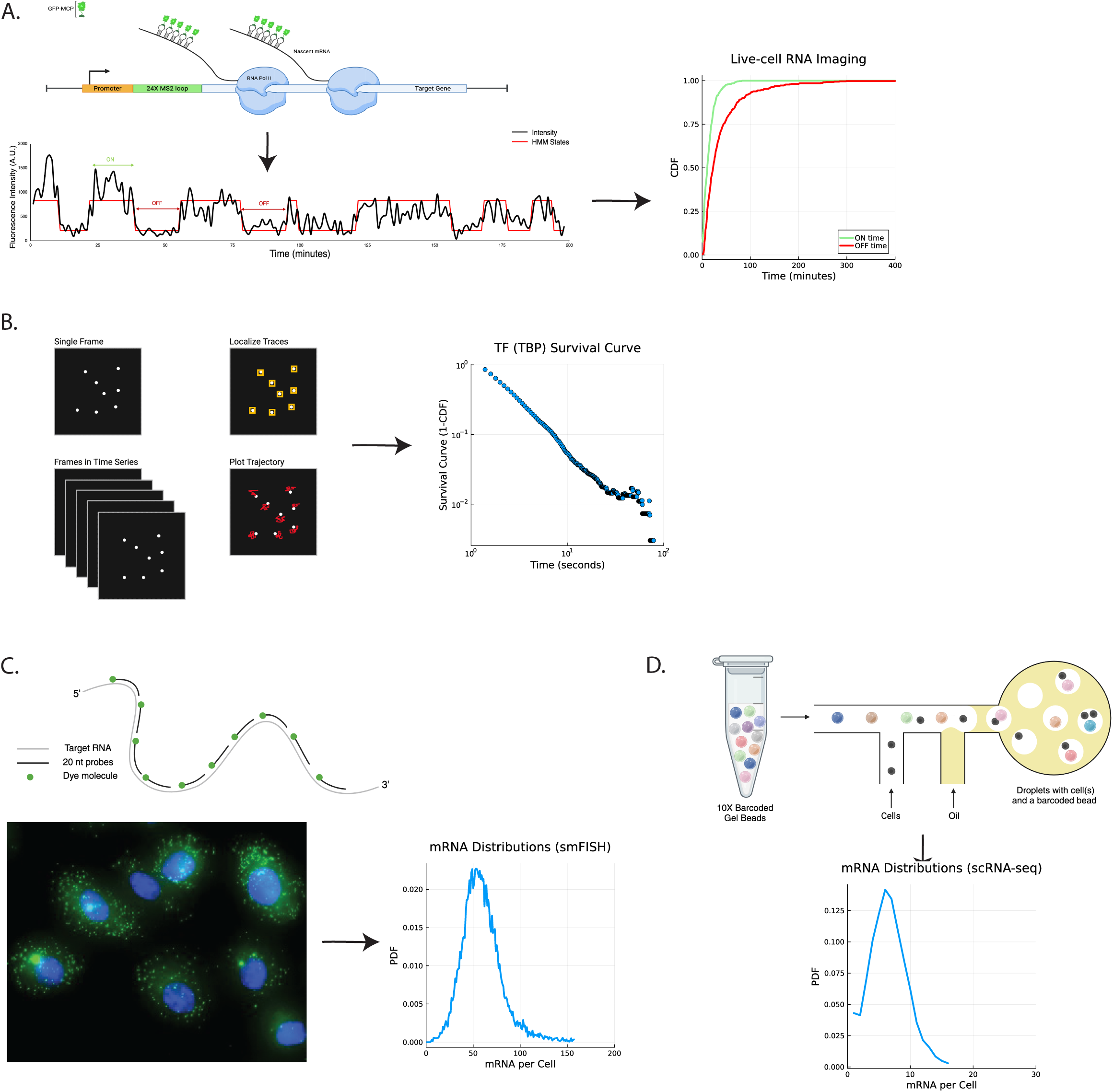
Methods for measuring single-cell gene expression. A) Left, illustration of 24XMS2 stem loop (green bar) fluorescent tagging of nascent mRNA of a target gene. MS2 stem loops are transcribed and tagged with GFP-MS2 coat protein (GFP-MCP) to enable visualization of nascent RNA at the transcription site (TS). Fluorescence intensity of TS is then measured and plotted in black and hidden Markov Model (HMM) fitting is used to identify ON and OFF time distributions in red. Right, Cumulative Distribution Function (CDF) for ON (green) and OFF (red) time distributions. B) Left, single-molecule tracking (SMT) of a TF of interest and its survival curve (1-CDF), JF646 dye was used to label the TF of interest, and 4-minute movie was acquired with 200ms interval. C) single molecule fluorescent *in situ* hybridization (smFISH) and its abundance distribution. smFISH probe sets were designed using Stellaris Probe Designer using Quasar 570 and 670 dyes. D) single cell RNA sequencing (scRNA-seq) and its abundance distribution. scRNA-Seq libraries were prepared using Chromium Single Cell 3′ Reagent Kits (10X Genomics) according to the manufacturer’s protocol. With double 10bp index cycles, 28 forward and 90 reverse cycles were run on NovaSeq6000 machine.

We integrated smFISH to count the number of transcripts in single cells. smFISH is a versatile method for analyzing single-cell gene expression, ranging from a few genes to highly multiplexed implementations [41–46]. Here, we focus on 30 genes across a range of expression levels, including the same ones used for live-cell imaging [21]. These datasets served as our best reference standard for absolute mRNA counts per cell, owing to the high detection efficiency of smFISH and hence the ability to provide both spatial and quantitative gene expression measurements (Figure 1C) [46, 47]. While advances in smFISH technologies have enabled precise visualization and enumeration of individual mRNA transcripts, offering critical insights into transcriptional regulation and cellular heterogeneity [46], smFISH remains limited in several respects. Most notably, it is restricted to fixed cells, precluding temporal resolution [48], and the time-intensive labeling/imaging approaches and cost typically constrain sample throughput.

To address these constraints on cost and scalability, we integrated scRNA-seq as an optional extension for quantifying transcripts in single cell resolution. While our primary approach integrates live-cell imaging, smFISH, and SMT to enable detailed mechanistic inference, scRNA-seq provides a complementary, transcriptome-wide view of gene expression at single-cell resolution (Figure 1D) [49–51]. This advantage makes it particularly useful in settings where the imaging techniques are not feasible at larger scale. In particular, scRNA-seq has dramatically advanced our ability to explore how gene expression varies across individual cells [52–54]. However, scRNA-seq has known limitations that affect its quantitative accuracy. A major challenge is transcript dropout, a prevalent technical artifact that results in zero counts which can obscure biological interpretation, as these zeros may reflect either true absence of expression or inefficient capture and reverse transcription [55–57]. Indeed, capture rates can be lower than 10% for many mRNAs [56]. For this reason, we treat exonic smFISH as the gold standard for absolute mRNA quantification in HBECs, while leveraging scRNA-seq for transcriptome-wide kinetic inference. In summary, our framework integrates complementary single-cell approaches spanning live-cell, fixed-cell, and sequencing-based measurements, balancing mechanistic resolution with scalability to generate a comprehensive dataset for downstream mathematical modeling.

We performed a comparative analysis of stochastic transcription models of increasing mechanistic complexity, aiming to determine which model most comprehensively captures key biochemical events during transcription, including elongation and splicing. All models are built on the canonical telegraph model, in which a gene switches stochastically between transcriptionally active (ON) and inactive (OFF) states (states are denoted by *G*). We extended this framework by explicitly modeling two additional processes, elongation and splicing. First, elongation is represented as a series of *R* discrete steps, each corresponding to the position of a nascent RNA molecule as it progresses along the gene body [21]. A nascent RNA advances to the next step only if that step is unoccupied, mimicking the fact that RNA polymerases cannot overtake one another. When the nascent RNA reaches the final step, it is released as a mature mRNA. Second, splicing is incorporated through S sites distributed along the elongation track, representing positions where co-transcriptional splicing can occur as the transcript is being synthesized. Each R step can be either occupied or unoccupied, and irreversible forward transitions between R steps occur only if the next R step is unoccupied. An mRNA molecule is ejected from the final R step. Splicing sites are locations (up to a maximum of R) where the mRNA can be potentially spliced. When the gene is in the active G state, there is the possibility that the first R step can be occupied if it is currently unoccupied.

Within this shared framework, we evaluated three model variants (Figure 2A). The 2-state telegraph model (G=2) captures basic ON/OFF gene switching [15]. The 3-state model adds an intermediate inactive state, allowing it to represent additional regulatory processes/transitions such as chromatin remodeling or pre-initiation complex assembly [19, 21, 58]. The kinetic proofreading (KP) model further extends the 3-state model by introducing an irreversible, energy-consuming transition from the ON state back to the initial OFF state, breaking detailed balance and enabling a form of regulatory specificity that thermodynamic equilibrium models cannot achieve (Figure 2A, top, middle, and bottom panels, respectively) [27].

**Figure 2.**
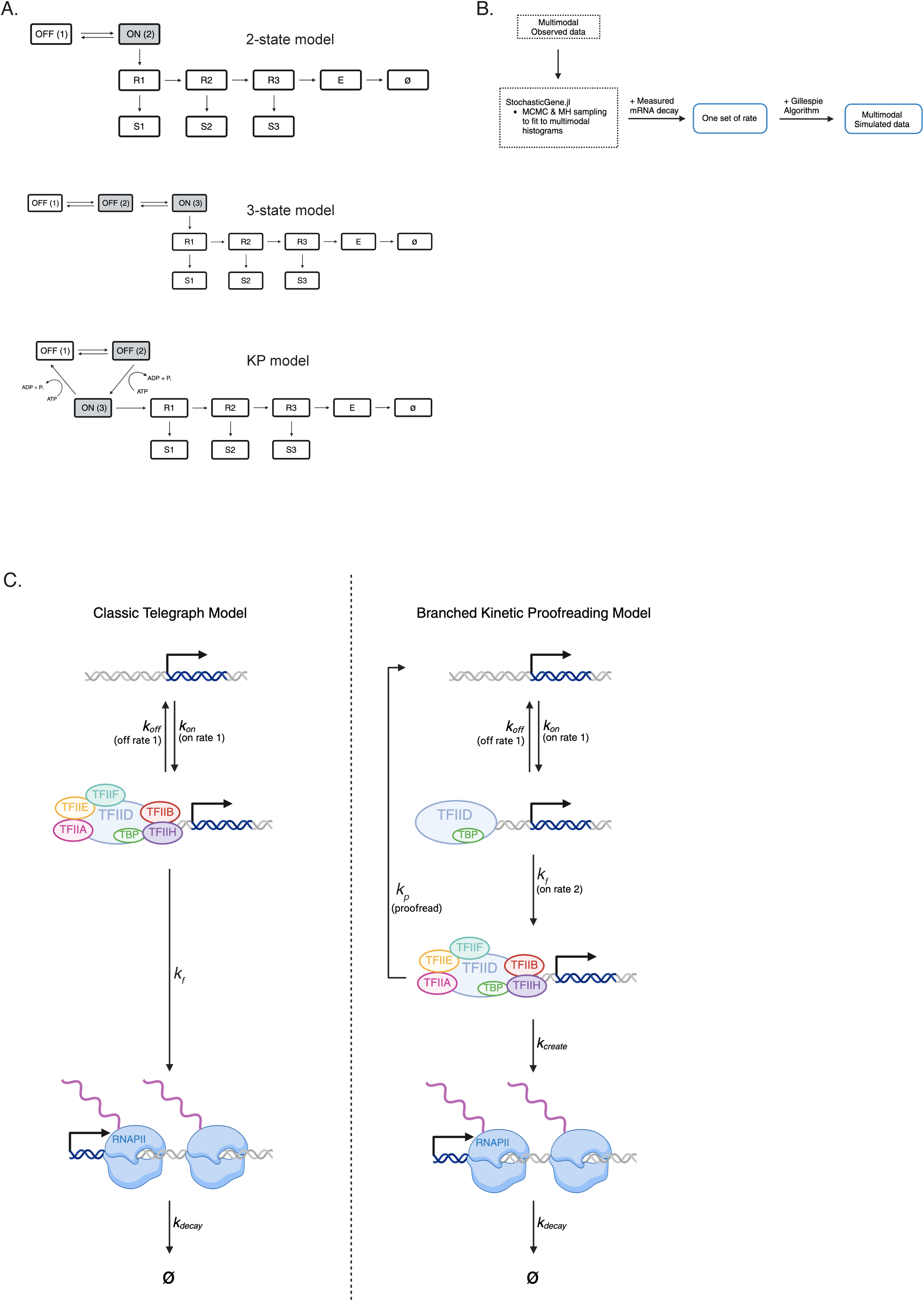
DiHerent transcription models. A) Illustration of 2-state, 3-state, and kinetic proofreading (KP) models. Grey boxes are TF binding states. R stands for RNA elongation, S stands for splicing, and E stands for mRNA eject. B) Schematic of rate inference algorithm. C) Comparison of classic telegraph model and branched kinetic proofreading model. Note that there is an extra step in KP model for enhanced specificity. *k_on_* is the first ON rate, representing the transition from the fully inactive state to the active state in the classic telegraph model, or to the intermediate state in the KP model. *k_off_* is the OFF rate that governs the transition from the active or intermediate state back to the fully inactive state. In the classic telegraph model, *k_f_* represents the mRNA production rate, whereas in the KP model, it represents the second ON rate corresponding to the transition from the intermediate state to the active state. *k_p_* is the proofreading rate that evaluates the specificity of the TF binding and returns the system to the inactive state if proofreading fails. Finally*, k_create_* denotes the mRNA production rate in the KP model.

With these models of transcription, we then sought to incorporate the dynamics of general TFs (GTFs). Although the molecular identities of individual gene states remain relatively ill defined, active states are strongly associated with GTF binding [39, 40]. Therefore, GTF dynamics are expected to influence transitions between gene states that differ in their levels of GTF occupancy. More specifically, in the 2-state model, there is only a single ON state which might correspond to TF/GTF binding (Figure 2A, top panel has only one gray box). In the 3-state model, the intermediate state can be interpreted as the GTF-bound PIC stage, accommodating upstream regulatory events such as TF and RNAPII recruitment. The KP model is similar to the 3-state model with the added possibility that the first OFF state represents potential nonspecific configurations, the intermediate state (second OFF state) represents GTF-bound configurations, and the ON state initiates transcription. There is an explicit energy-consuming process controlling GTF occupancy, which is essential to the increased discrimination between specific and non-specific binding [59]. As depicted in Figure 2A, the gray boxes correspond to a TF/GTF bound state. The distribution of bound times is experimentally determined by single-molecule imaging in living cells and reported as a TF survival curve, which is the relative frequency of a bound state, or (1-CDF), where CDF is the cumulative distribution function of dwell times. Intuitively, one can start a timer upon TF/GTF binding (corresponding to occupancy of a gray gene state) and stop it upon unbinding (corresponding to occupancy of a white gene state), yielding the dwell time distribution. To further constrain these models, we incorporated GTF dwell times measured by SMT, alongside previously used datasets such as steady-state mRNA distributions and transcriptional bursting kinetics from live-cell imaging [37, 60–64]. This additional modality improves parameter identifiability and biological realism. In particular, we focused on TBP, a core subunit of TFIID central to promoter recognition and PIC assembly (Figure 2B, C) [39, 40, 65]. TBP dwell times have been measured in multiple studies [66, 67] and correlate with transcriptional activation in HBEC [38], making them a natural choice for linking GTF dynamics to transcriptional state transitions. Overall, we focus here on a set of biologically plausible scenarios by jointly analyzing smFISH, nascent mRNA live-cell imaging, and GTF dwell time datasets for the ten aforementioned genes, using TBP as the representative GTF (Figure 2C). Posterior distributions of the fitted rates reflect excellent convergence of the rates by showing a distinguishable peak, indicating convergence in the likelihood estimate (Figure S1). These sharp posterior distributions reflect strong agreement between the model and input data, bolstered by informative priors such as multi-modal dataset distributions and multiple experimental constraints.

To highlight the differences between the models we focused on two representative genes: *ERRFI1* and *RHOA* (Figures 3A and 3B, respectively). For *ERRFI1*, although all models reproduced the mRNA and burst duration distributions well, the 2-state model overestimated GTF dwell time due to unrealistically slow on and off rates (0.012 and 0.043 per minute, respectively). These results are consistent with the 2-state model being too coarse to recapitulate complex bursting behavior or multi-step regulation in all cases, as has been previously demonstrated [19, 21]. The 3-state and KP models more accurately captured all distributional features, reinforcing the necessity of intermediate gene states for this gene. Indeed, best fit models, 3-state and KP models, show relatively equal occupation of three distinct gene states (Figure 3A). In contrast, *RHOA*, which could be classified as either a 2 or a 3-gene state gene, demonstrated ambiguity by showing that all three models recapitulated most features, including TF dwell time distribution. However, the mRNA distribution of *RHOA* in the 2-state model did not replicate the experimental data as accurately as the 3-gene state models (Figure 3B). Furthermore, the OFF-time distribution for the gene *RHOA* in the 2-state model also failed to match the experimental distribution as effectively as the 3-gene state models. This observation led us to conclude that despite infrequent transitions of *RHOA* to the third gene state (due to low ON rate 2 and proofreading rate), inclusion of a third state improved alignment of simulated OFF time distribution, suggesting that incorporating the third gene state, even as a placeholder, is essential for accurately recapitulating the kinetic distributions.

**Figure 3.**
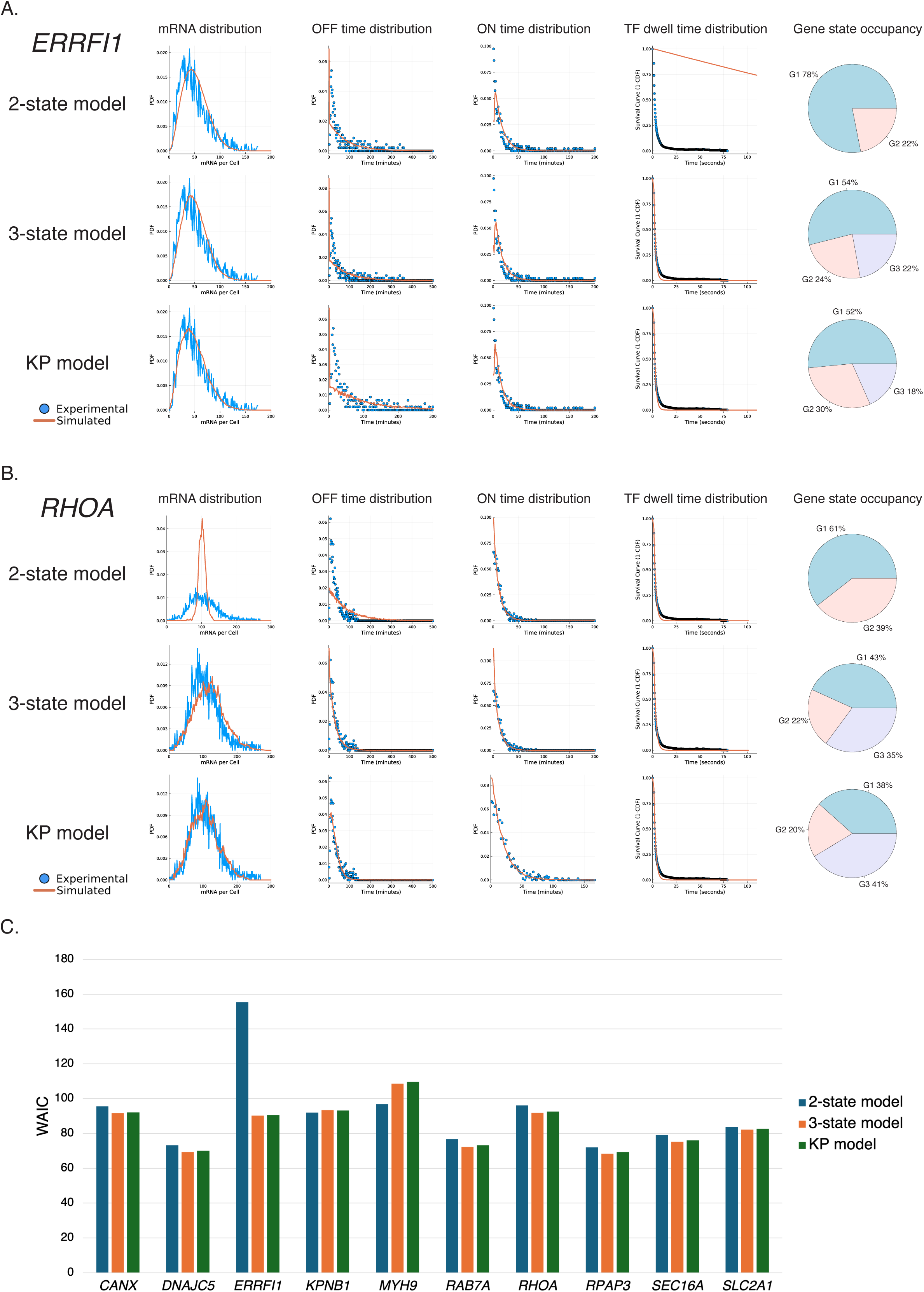
Comparison of distributions and gene occupancy pie chart. A) *ERRFI1* and B) *RHOA* across all 3 models (2-state, 3-state, and KP). Blue lines and dots indicate empirical distributions from different measurements (mRNA distribution, live-cell imaging for nascent mRNA, and SMT for TF dwell time), and orange line represents simulated distributions from the model using inferred rates. The pie charts indicate the gene state occupancy based on the inferred rates to explore how much time this system spends in each gene state. The genes are simulated until convergence, whichis explained in detail in Methods. C) Watanabe-Akaike Information Criterion (WAIC) measurement for 10 genes with standard error bars. Lower WAIC indicates better balance of a model.

To evaluate alternative models, we first inferred transcriptional rate parameters using a previously published stochastic modeling framework (StochasticGene.jl) based on Metropolis-Hastings MCMC sampling [19, 21]. Upon convergence, the algorithm yielded a set of kinetic rate parameters and posterior likelihood distribution that best reproduced the input distributions derived from the experimental data (see Methods). On top of the inferred rate, the StochasticGene.jl workflow also delivers Watanabe-Akaike Information Criterion (WAIC) (Figure 3C) [32], a type of Bayesian approach designed to evaluate and compare models by balancing their complexity with their likelihood. Lower WAIC value indicates that a model has better balance between fit and complexity; therefore, it is likely to produce better inference. From the WAIC measurement, *ERRFI1* exhibited the strongest preference for the 3-state and KP model, with ΔWAIC of -41.38 ± 0.12 relative to 2-state model, respectively. For most other genes, the 3-state and KP models yielded similar WAIC values, both slightly outperforming the 2-state model and suggesting the presence of intermediate gene states not captured by simpler models. Only one gene (*MYH9*) unambiguously showed better performance for a 2-state model. In summary, we can use WAIC-guided model selection to identify parameter combinations which effectively describe both dynamic and static measurements of gene expression in single cells. Importantly, this integrative, multimodal framework enables us to discriminate between competing mechanistic models that would otherwise appear equivalent when evaluated using any single data modality alone, thereby providing a more stringent and biologically grounded basis for inferring transcriptional regulatory mechanisms.

### Kinetic proofreading as a general model for gene expression data

Our ultimate goal is to use mathematical models for biochemical inference, and one important consideration that is not captured by treating genes in isolation is how TFs quantitatively discriminate one gene from another [68]. In this section, we develop a quantitative framework to evaluate transcriptional specificity across gene expression models. We first formalize specificity using a kinetic competition metric that distinguishes transcription driven by specific versus non-specific TF binding (Figure S2). Using this definition, we analyze how specificity emerges in the 2, 3-state and KP models under biologically realistic assumptions, including excess non-specific TFs and modest differences in binding affinities. We then derive analytical expressions for each model and compare their limits, highlighting the conditions under which additional intermediate states and energy-dependent steps enhance discrimination. Finally, we use this framework to motivate the adoption of the KP model for subsequent analyses.

The 3-state model and KP model performed comparably in terms of fitting experimental data, and we now explore the conceptual advantages and disadvantages of these models in terms of specificity. Both models have multiple OFF states where activator identity can be sensed, for example by having more than one step in the activation pathway reflecting TF identity. Both models can be made into non-equilibrium models which break detailed balance. Historically, the branched checkpoint in the KP model is assumed to be necessary as a critical regulatory mechanism that is not present in the 2 or 3-state telegraph models [27, 69](Figure 2A). However, we show below that branched models are not explicitly required for proofreading in some rate regimes.

In general, the KP and 3-state models have the capability to extend beyond equilibrium by introducing irreversible or biased, ATP-driven steps which break detailed balance. The inclusion of an extra ATP-driven step improves specificity by reducing the likelihood that nonspecific (or unintended) GTF binding leads to productive transcription (Figure 2B) [27, 69]. Previous work has demonstrated, for example, that ATP-consuming processes such as chaperone, proteasome, and chromatin remodeling activity directly shape both gene activation and TF dynamics in living cells [65]. This accommodation is particularly important given the cellular milieu, where nonspecific TFs are often present in excess relative to their specific counterparts [59]. Given that TF binding to DNA occurs via diffusion, higher concentration of nonspecific TFs increases the probability of encountering the DNA binding site [70]. Thus, specificity likely depends primarily on off rates: correct TFs typically dissociate more slowly than incorrect ones, but this difference alone yields only modest discrimination. For instance, human specific TFs typically recognize DNA motifs of 6-20 bp [70], reviewed in [71], yet the difference in binding energy between specific and nonspecific interactions is only about an order of magnitude [72, 73], suggesting that in the 2-state model correct and erroneous binding events can occur at comparable frequencies. By introducing a second, energy-driven checkpoint, the 3-state and KP models effectively reads TF identity twice: once at initial binding and again during the intermediate state. This double discrimination sharply amplifies fidelity [59]. Formally, specificity in the proofreading scheme can approach unity because both dissociation and proofreading rates contribute multiplicatively to the probability of rejecting incorrect substrates [72].

To formalize these concepts, we define the concept of transcription specificity using a kinetic competition framework which is uniquely suited to single-molecule experimental approaches (Methods). Consider a gene that is activated in response to the binding of a specific TF to a cognate binding motif located, for example, in the promoter or an enhancer. A non-specific factor may also bind this motif, but with a higher off-rate and hence lower affinity. These TFs are examples of what has previously been labeled single “target factors” [74]. This non-specific transcription activator can also initiate downstream steps of transcription and could indeed function as a specific activator at another gene. As such, the TFs interact with or “recruit” GTFs or what has previously been called “multitarget factors,” which work at many genes. We define the quantity *ψ* as:

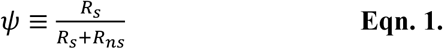

Where *R_s_* is the number of RNA transcripts produced due to the binding of the specific transcription factor, and *R_ns_* is the number of RNA transcripts produced due to the binding of the non-specific transcription factor. Here *ψ* is motivated by the same quantity in the analysis of RNA splicing, which is the “percent spliced in” (p.s.i) of a certain isoform. *ψ* ∼ 1 corresponds to high specificity; *ψ* << 1 is low specificity. Using this defined quantity with the different models described above allows us to explore the limits of specificity with a gene-centric approach.

We first look at the classic 2-state telegraph model (Figure 2C), but in the competition framework (Figure S2). In the competition model, the rates correspond to the same steps in the single-gene activation model but now with subscripts designating specific (*s*) or non-specific (*ns*). Suppose *k_on_s_* is ON rate for specific (correct) binding of TF and *k_on_ns_* is ON rate for a single nonspecific TF (incorrect). Similarly, if we define *k_off_s_* and *k_off_ns_*, the result for specificity would then be as follows:

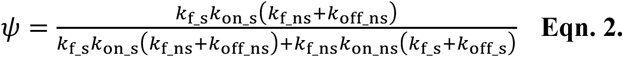

Note that the *k_f_* terms appear explicitly in the final result owing to the particular topology of the telegraph model. Intuitively, this dependence is due to the fact that if *k_f_* were extremely fast, any random encounter between the TF and DNA would lead to transcription initiation, and thus binding events become locked in. In fact, if we compare two different TF present at equal abundance and assume that ON rates for the specific and non-specific factors are the same, which is a reasonable assumption (albeit experimentally untested in living cells to our knowledge), and *k_f_* is both fast compared to binding dynamics but also similar for the specific and non-specific TF (since both TFs are capable of leading to transcriptional activation and differ only in DNA binding), then an asymptotic expansion for *k_f_* >>1 gives the following result:

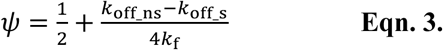

Now the specificity is ∼ ½ to first order and improves proportionally with the difference between non-specific and specific OFF rates with respect to the forward rate, which is large in this instance. The net result is little specificity, again assuming that both specific and non-specific factors lead to transcriptional activation with similar likelihood.

Next, we turn to the scenario most likely to capture the cellular conditions. First, all available data indicates that *k_f_* is much slower than the *k_on_* and *k_off_*, which is the reason that transcription is inefficient [75] and bursting infrequent in human cells [21]. Second, there is not just one non-specific transcription factor but hundreds or more, which we mathematically capture by increasing *k_on_ns_*. For the sake of simplicity, we assume that the non-specific ON rate is *M* times greater than the specific ON rate to account for the higher concentration of the non-specific factors, and we also assume the non-specific factors bind *M* times weaker, i.e. *k_off_ns_* = *M*\**k_off_s_*. This latter assumption is motivated by measurements that indicate non-specific TF binding is more transient than specific binding, with dissociation rates differing by 10- to 100-fold [37, 67]. The asymptotic result gives:

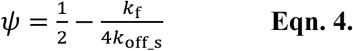

where we assume M>>1. Now, the specificity has a *maximum* value of ½, which is only achieved in the limit of slow forward progression (*k_f_* << *k_off_*). Thus, using the telegraph model under conditions which approximate intracellular conditions results in a problem of specificity. The root of this problem is captured in the mathematical assumption made here—based on experimental data—that there are many non-specific factors yet the difference in OFF rate between specific and non-specific factors is simply not great enough to provide contrast based on this simple 2-state model. This argument has been made previously based on existing experimental data for specific and non-specific TF dwell times measured by SMT and also proteomic studies of the number of TF per cell [19, 21, 37, 67]. Our goal then is to see if alternate models, constrained by these same experimental realities, provide greater specificity.

The primary advantage of 3-state models is the chance for another OFF rate from the intermediate state which can be activator-specific. In the model topology we consider here, the linear 3-state model includes this additional OFF rate as a backward rate from ON (3) to OFF (2), and the KP model contains an additional rate from ON (3) to OFF (1) (Figure 2A). These backward rates can be set lower for the specific activator compared to the non-specific activator, thus allowing the identity of the TF to be ‘sensed twice’, which is one condition of a proofreading scheme as originally described [27, 69]. The other condition—that the second OFF rate involves a branched or cyclic pathway—is met in the KP topology but not in the other linear 3-state topology. In our KP model, non-specific complexes are completely disassembled, bringing the gene back to the OFF (1) state. Biochemically, one might consider any process capable of removing transcription complexes from chromatin, which might be chromatin remodelers [76, 77], chaperones [78] or the proteasome [79, 80]. Note that we do not examine any cases where ON rates can also show specificity (see Discussion). Here, we now take a general look at how these 3-state models potentially influence specificity under a limited set of assumptions.

The equation for specificity for the KP and linear 3-state models under the intracellular assumptions, respectively are (see also Methods):

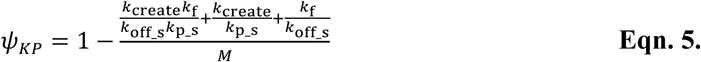

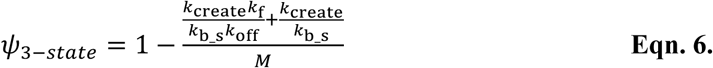

Now *k_f_* is the transition rate to the intermediate state, and *k_create_* is the transition from the intermediate state to active RNA synthesis. The above equations come from keeping first order terms in the expansion around *k_f_* and *k_create_* going toward zero, which corresponds to the assumption that the forward rates are small compared to the backward rates, as described above. *k_p_* and *k_b_* are present in models with more than 2 states. *M* is again defined for simplicity to be the ratio between specific and non-specific ON rates *and* specific and non-specific OFF rates (*k_on_ns_ = M*k_on_s_; k_off_ns_ = M*k_off_s_; k_b_ns_ = M*k_b_s_; k_p_ns_ = M*k_p_s_*). At the molecular level, this scenario would correspond to the case where there are *M*-fold more non-specific TFs than specific ones, but their dwell time on chromatin is *M*-fold lower, and the non-specific transcription complexes are *M*-times less stable. Again, specificity is thus sensed twice in these models: first at initial binding and then after recruitment of downstream effectors, the only difference being whether the backward rate is to an intermediate occupancy state or a fully unoccupied state.

The difference in specificity for these models can be computed by subtracting these two quantities under conditions where *k_b_* = *k_p_*≡ *k_off_int_*. Thus, the backward rates are the same and generically referred to as the intermediate off rate, *k_off_int_*, and all other rates are the same, and only the relative topology is assessed. The result is:

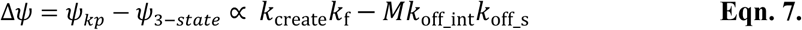

Indicating that the relative improvement in specificity for each model is rate dependent. Intuitively, one can see that if forward rates dominate (*k_f_, k_create_* > *k_off_s_, k_off_int_*), the KP topology outperforms the linear 3-state topology in specificity, because there is an advantage in forcing the gene back to the unoccupied state, i.e. driving complexes off of chromatin. This regime is the classic Hopfield proofreading regime: the build-up of intermediate states which could potentially drive forward progress through mass action is prevented. If however, the backward rates (*k_off_int_*, *k_off_s_*) are large, then the branched KP model provides no benefit because the backward flux almost always ends in the fully unoccupied state anyway, without the buildup of intermediate states.

For either model topology, it is clear that the maximum specificity is now unity. These models break detailed balance and therefore require energy input, but the net result is that TF identity is determined at multiple steps and many binding events are therefore non-productive. Thus, the addition of an intermediate state which introduces a delay between initial TF binding and transcriptional activation can lead to arbitrarily high levels of discrimination between specific and non-specific TF binding, but at the cost of efficiency.

Without loss of generality, we default to the branched KP model for subsequent data analysis in this study for several reasons. First, multi-state models provide a quantitative improvement for nearly all genes analyzed and a definitive improvement for at least one gene (*ERRFI1*), and these models do not limit analysis of other genes (Figure 3). Second, the 2-state telegraph model suffers from a conceptual dilemma not captured in single gene analysis, which is the inability to sufficiently discriminate specific and non-specific binding, as described above. Third, the linear 3-state and branched KP models are nearly indistinguishable under the relaxed assumptions used here, but the KP model has the benefit of applicability in other systems, previous use for gene regulation in single cells [28, 80–83], and better alignment with biochemical mechanisms (see Discussion). Nevertheless, we hasten to add that because the models are so similar, the conclusions that are reached with the branched KP model likely apply to the linear 3-state model under the assumptions used here. In conclusion, we therefore adopted the KP model for subsequent genome-wide inference of kinetic parameters, leveraging its ability to integrate multiple experimental modalities and to represent transcriptional control with greater biological accuracy.

The complete set of rate parameters inferred for the ten genes for the KP model is in Table 1. These rates are: two activation rates (ON rate 1, *k_on_*; ON rate 2, *k_f_*), two deactivation rates (OFF rate 1, *k_off_*; Proofread, *k_p_*), elongation and splicing rates (note that in the simulation step there are three elongation and three splicing, each with same rate), an eject rate, and a transcript decay rate. In the full stochastic simulation, *k_create_* (defined above for the ordinary differential equation competition model) does not have a 1:1 analog in the stochastic model, since the stochastic model also describes the details of elongation and splicing. Depending on the experimental design and model configuration, the eject rate can represent either mRNA synthesis or ejection of the reporter gene. Simulated data generated from these parameters closely recapitulated the empirical distributions of mRNA per cell, ON/OFF durations, and GTF dwell times across all genes (Figure S2).

**Table 1.**
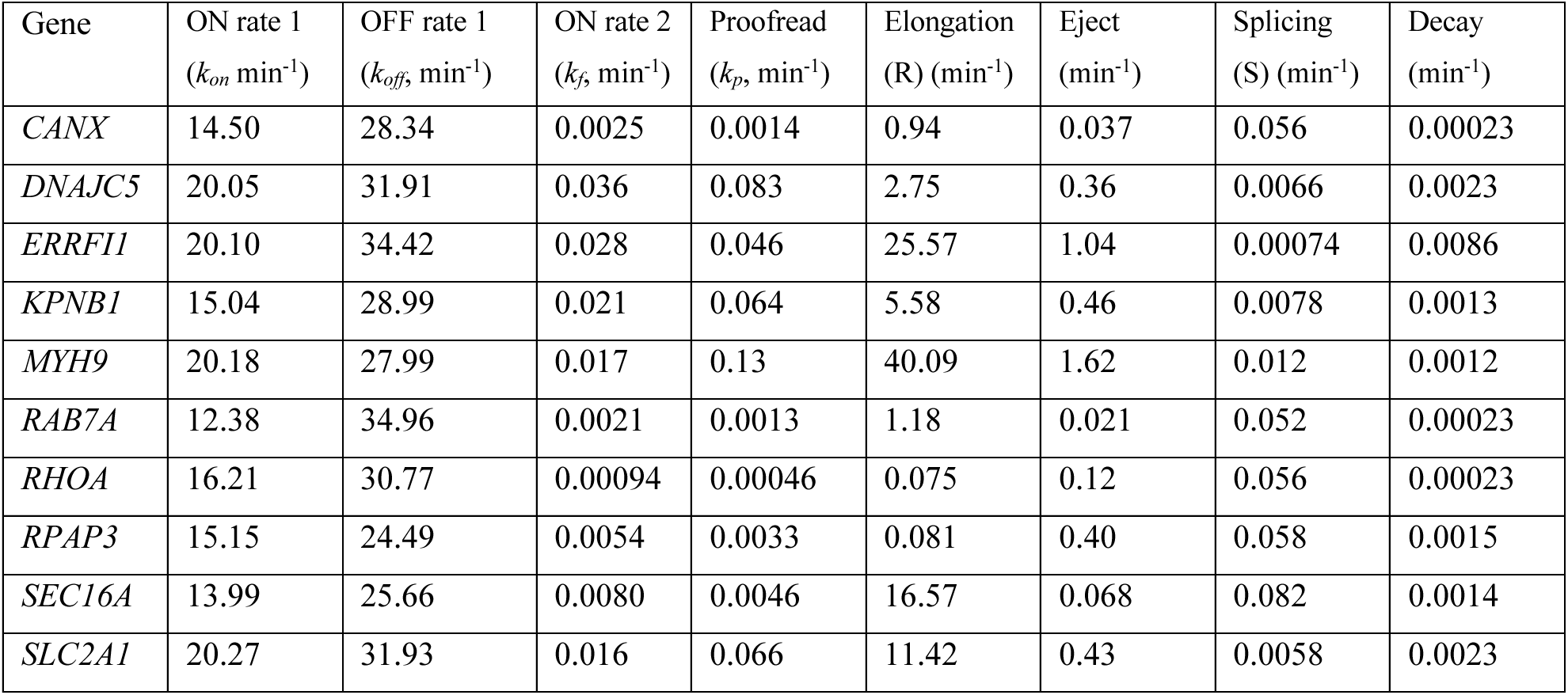
List of rates of 10 genes using smFISH, nascent mRNA live-cell imaging and GTF (TBP) dwell time live-cell imaging as inputs.

There are several findings of note in the inferred kinetic rates in the KP model (Table 1). First, ON rate 1 and OFF rate 1 were relatively invariant; the range of the ON rate 1 is roughly from 12 to 20 min^-1^, and the range of the OFF rate 1 is roughly from 24 to 35 min^-1^. These values are tightly constrained by TBP survival curves, leading to an OFF rate which we experimentally measured at ∼14 min^-1^. Second, we observed that elongation, transcript creation, and splicing rates are compensatory: when one is fast, the other parameters tend to be slower, suggesting identifiability challenges in resolving these contributions independently. Given that mammalian splicing durations vary widely depending on intron length and gene context, with prior studies estimating ∼1 to 40 minutes of endogenous gene introns or small intron splicing[81], we imposed a tight prior on the splicing rate to be 20 minutes (0.05 min^-1^), which will adjust the rate-limiting step and the ON time distributions.

Importantly, we observed that several parameter ratios appeared to be conserved. We therefore defined two dimensionless constants to quantify transcriptional regulation: the ratio between ON rate 1 and OFF rate 1, which we called the search constant (*K_search_*) and the ratio between ON rate 2 and proofread rate, which we called the proofreading constant (*K_proof_*). These ratios are equivalent to equilibrium molecular association constants (*K_A_*) and capture the relative tendency of transcriptional activation to be favored through a bias toward increased search efficiency or decreased proofreading. From the smFISH dataset, the median of *K_search_* was 0.58 and the median of *K_proof_* was 1.18. Notably, these ratios remained relatively stable: 0.47 and 0.92 for *K_search_* and *K_proof_*, respectively, even after setting a tight prior on the splicing rate, indicating that key regulatory features are robust to constraints on downstream processing rates (Supplementary Table 1). We thus propose these constants as dimensionless shape parameters for fitting single-cell expression distributions, much in the same way that ‘burst size’ has been used in many previous studies [18, 22, 31]. The conservation suggests that cells maintain a consistent regulatory balance across genes: early binding is dominated by dissociation (*K_search_* < 1), while the intermediate state is poised for progression or disassembly (*K_proof_* ∼ 1).

In subsequent analyses, we compare *K_search_* and *K_proof_* inferred directly from scRNA-seq dataset to investigate their conservation across modalities and genomic scale.

### Extension of model simulation to scRNA-seq dataset gives genome-wide insight to the gene regulatory mechanism

We next aimed to assess whether scRNA-seq measurements could be used to infer model parameters, even in the absence of nascent RNA live-cell imaging or smFISH. Our approach was to use the detailed analysis based on comprehensive characterization of our 10 gene training datasets to establish strong priors for analysis of scRNA-seq distributions. Additionally, we aimed to incorporate TBP dwell times to constrain the fit to scRNA-seq distributions. Although scRNA-seq has the potential for transcriptome-wide coverage of gene expression heterogeneity, the technology suffers from technical noise, including low capture efficiency and transcript dropout, which can lead to underestimation of true mRNA abundances [84]. Therefore, any quantitative model needs to account for this feature.

We first needed to calibrate our scRNA-seq counts against those from smFISH dataset, with the goal of estimating a global yield factor for our model to account for the capture efficiency. After quality control and filtering of the scRNA-seq dataset (Figure 4A; see Methods), we next selected 30 genes for smFISH, which we carried out in the same cell line under identical culture conditions. Plotting the mean expression values revealed a systematic undercounting in scRNA-seq relative to smFISH (Figure 4B). We performed reduced major axis regression constrained to pass through the origin, thereby assuming that zero expression in smFISH should correspond to zero in scRNA-seq. The slope of the fitted line, representing the yield factor, was 0.068 ±0.0131, indicating that, on average, fewer than 1 in 10 mRNAs detected by smFISH are captured in scRNA-seq under our experimental conditions. We further examined this relationship at the level of individual genes by plotting per-gene scatter plots, in which the red tick indicates the median across all bins for a given gene, providing a single-number summary of the deviation of scRNA-seq counts from the ground-truth smFISH counts.

**Figure 4.**
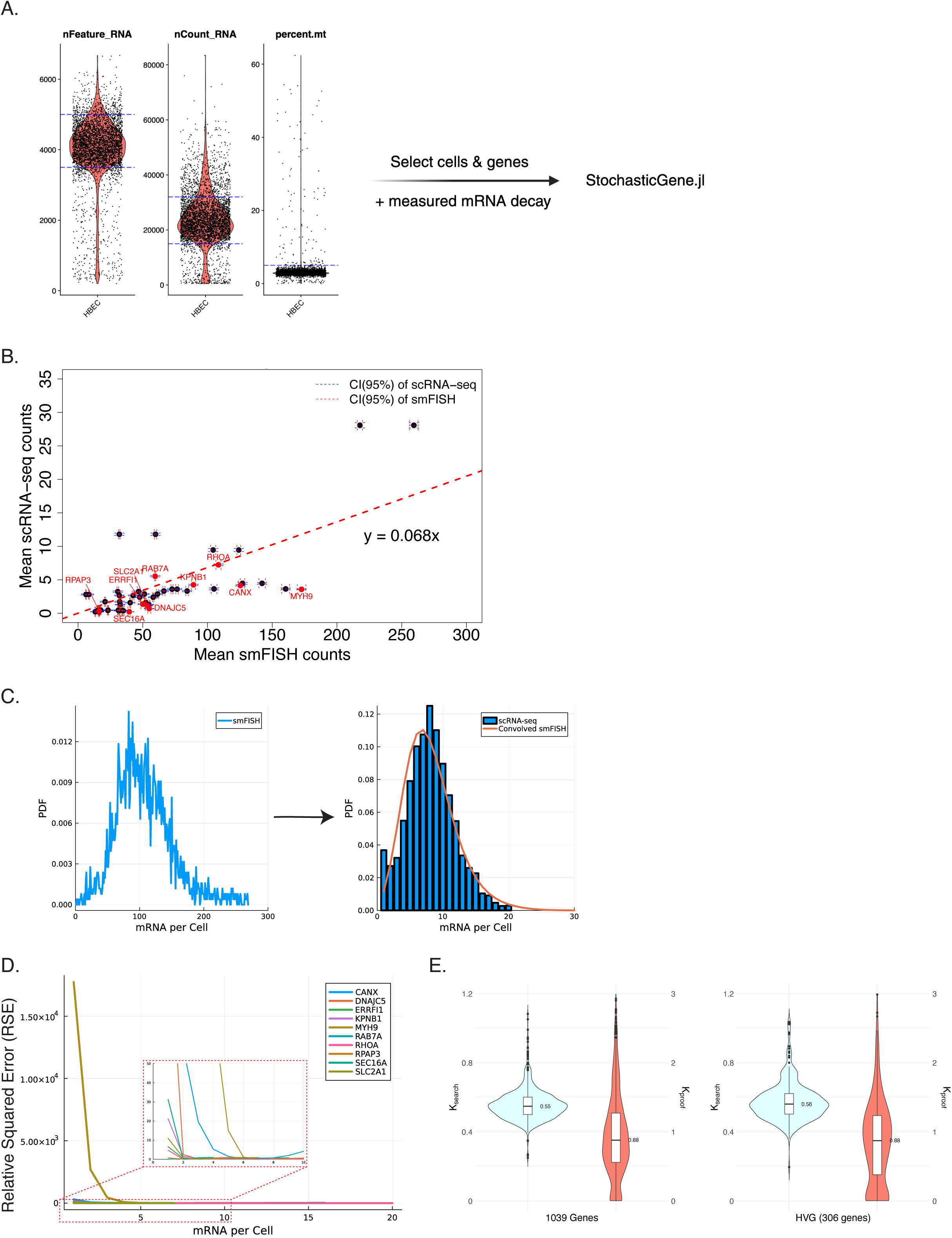
Comparison between scRNA-seq vs. smFISH datasets. A) Quality control filtering of the scRNA-seq dataset. Cells with 3,500-5,000 detected genes in each cell (nFeature_RNA), total of 15,000-32,000 detected molecules within a cell (nCount_RNA), and a mitochondrial transcript fraction (percent.mt) below 5% were retained for stochastic gene simulations. Low nFeature_RNA may indicate dead, dying, or empty droplet, high nCount_RNA may indicate a possible multiplet. B) Scatter plot comparing raw scRNA-seq and smFISH measurements. The slope (0.068) of the reduced major axis regression with the line constrained to go through zero was used as a global yield factor to account for scRNA-seq dropout. Error bars for each gene indicate 95% confidence interval (CI) of scRNA-seq (vertical) and smFISH (horizontal). C) mRNA count distributions from smFISH (left) and scRNA-seq (right). The scRNA-seq distribution is compared with the smFISH distribution convolved using the global yield factor (orange line in right panel). D) Mean Squared Displacement (MSD) analysis for the 10 genes, illustrating the impact of dropout in the scRNA-seq dataset. Inset: magnified view of the highlighted region. E) *K_search_* and *K_proof_* of TF of interest. 1,039 genes and 309 highly variable genes were selected from HBEC scRNA-seq 10X dataset.

This yield factor enabled convolution, converting smFISH counts to scRNA-seq-like distributions. For convolution, we modeled capture efficiency by applying a binomial sampling process to the smFISH mRNA counts, using the yield factor as the binomial success probability, as done previously [85, 86]. For example, for *RHOA*, which lies close to the regression line, the convolved distribution of smFISH counts (orange in Figure 4C) closely matched the empirical scRNA-seq distribution (blue bars in Figure 4C). Deconvolution, however, presents greater challenges due to the ambiguous origin of zero counts and increased sparsity in scRNA-seq datasets [55, 56, 87, 88]. To quantify the gene-level differences between scRNA-seq and convolved smFISH measurements, we computed relative squared error (RSE) as a function of mRNA/cell defined as:

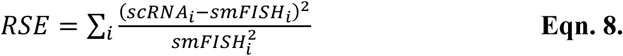

For each bin *i*, RSE captures relative deviation of scRNA-seq from the convolved smFISH (ground truth of our mRNA abundance), normalized by the smFISH expression level. In Figure 4D, RSE is plotted for each gene per bin counts in the mRNA abundance histogram (as shown in Figure 4C, right panel). We observed a sharp RSE decrease with increasing mRNA per cell, which suggests that low expressions are more prone to error in scRNA-seq, consistent with known dropout effects and quantification limits [55–57].

Using the empirically derived yield factor (0.068) and the kinetic proofreading model developed above, we fit scRNA-seq distributions from HBECs. Importantly, we constrained the fitting using GTF (TBP) dwell times. We further cross-referenced this set with decay rate measurements from HCT116 cells (12,872 genes) [87], and promoter motif annotations from the Eukaryotic Promoter Database (EPD) [89]. This filtering yielded 1,408 genes with TATA box motifs in the HBEC dataset, of which 1,039 genes had both decay rates and sufficient expression levels for kinetic modeling.

For each of these 1,039 genes, we inferred kinetic parameters using scRNA-seq mRNA distributions and TBP dwell time data and observed excellent recovery of the training set parameters. As in the smFISH analysis, the first two rates (ON rate 1 and OFF rate 1) were constrained by TBP residence time, while splicing rates were fixed near 0.05 min^-1^ to reflect biologically realistic values. Although ON/OFF time distributions were of course unavailable for each gene in this dataset, the fitted models successfully recapitulated experimental ON time distributions across the 10 genes where we did have live-cell data (Figure S3). We believe a central reason for this agreement was constraining the splicing rate—which largely determines the duration of fluorescent signals from active transcription sites—allowing us to accurately reproduce the shape and timing of the ON time distributions. We then computed the shape parameters, *K_search_* and *K_proof_*, from the underlying rates (Table 2) for the same set of 10 genes. Again, the rates were inferred only from scRNA-seq distributions and TBP dwell-time data. Despite methodological differences, the medians of *K_search_* and *K_proof_* values from the scRNA-seq-based inference (0.56 and 1.01, respectively) were remarkably consistent with those from the full dataset using smFISH and live-cell nascent RNA imaging (*K_search_*: 0.58 and *K_proof_*: 1.18). This agreement indicates that key aspects of transcriptional regulations, including GTF binding and proofreading kinetics, are robustly inferred across platforms.

**Table 2.**
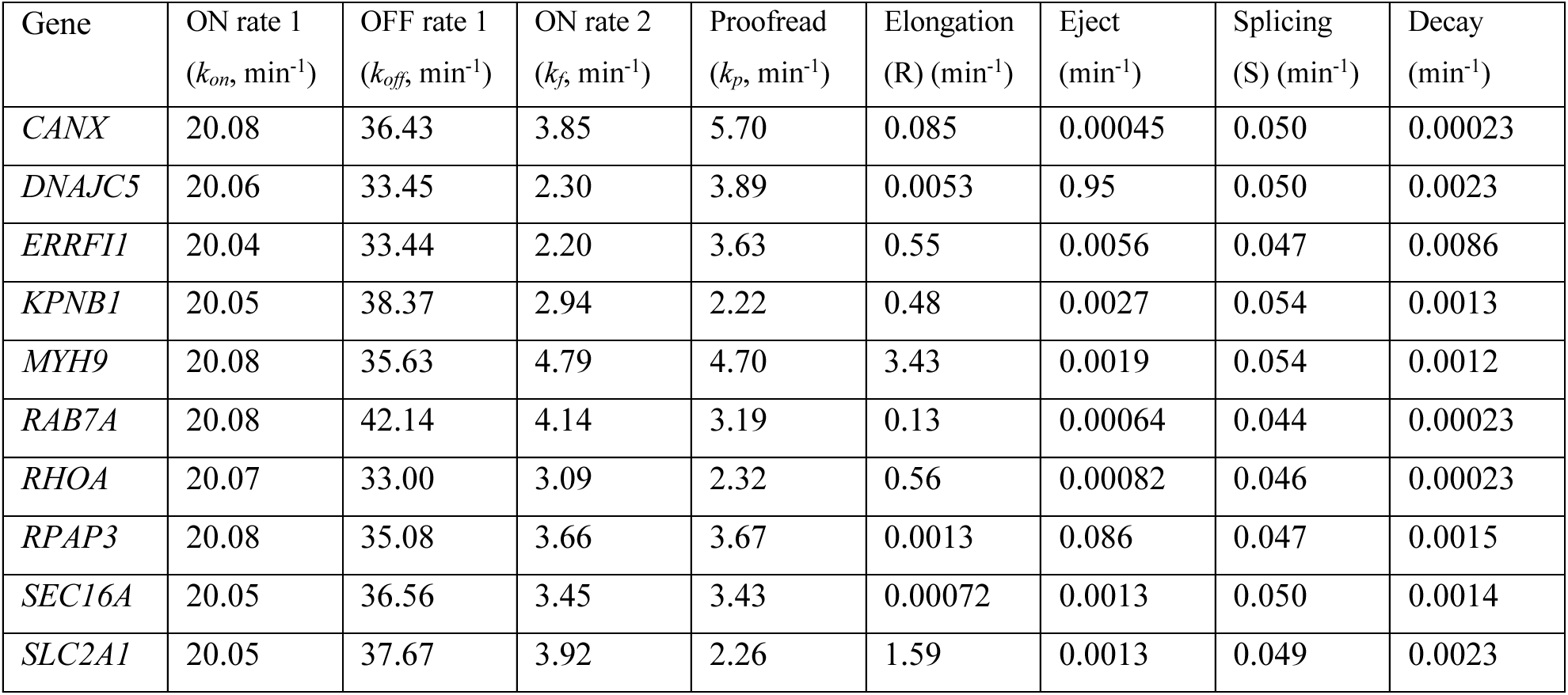
List of rates of 10 genes using scRNA-seq and GTF (TBP) dwell time live-cell imaging as inputs.

Expanding this analysis genome-wide, we computed *K_search_* and *K_proof_* values across all 1,039 genes, as well as a subset of 306 highly variable genes (HVGs) identified using the Seurat R package [90]. Among the 1,039 genes, the median of *K_search_* remained at 0.55, and the median of *K_proof_* was slightly lower at 0.88 (Figure 4E). Similar trends were observed in the HVG subset (*K_search_*: 0.56 and *K_proof_*: 0.88). As an additional control, we analyzed HVGs lacking TBP motif annotations. These genes also showed a conserved *K_search_* median of 0.55, but the *K_proof_* median was slightly higher (0.93), suggesting that promoter architecture influences proofreading dynamics. Taken together, these results suggest that while *K_search_* is a robust feature of transcriptional regulation under the constraints of our model, *K_proof_* may be more sensitive to promoter context or regulatory complexity. Notably, for a subset of genes, *K_proof_* approaches zero, indicating that proofreading strongly dominates over forward progression. In this regime, the KP model deviates most strongly from the 3-state model, highlighting the importance of the additional proofreading step. Conversely, as *K_proof_* increases toward large values, the KP model effectively reduces to a 3-state model, consistent with diminished influence of the proofreading pathway. These observations further support the use of the KP framework, as it captures both limiting behaviors and provides a more general description of transcriptional regulation. As such, *K_proof_* may be more informative for inferring transcriptional mechanisms from scRNA-seq data. These unitless constants thus offer a compact summary of the kinetic regimes governing TF search and commitment and may serve as useful parameters for characterizing transcriptional heterogeneity across the genome.

### The model predicts the action of individual TFs based on high throughput scRNA-seq screening

We hypothesized that changes in the gene expression distribution for individual genes might be used in concert with our quantitative model to infer basic mechanisms of transcriptional activation. As such, we next applied the KP model to a recent scRNA-seq ‘TF Atlas’ dataset to evaluate how overexpression of individual TFs modulates transcriptional kinetics, particularly the *K_search_* and *K_proof_* [33]. The TF Atlas study employed a multiplexed overexpression of regulatory factors (MORF), leveraging a barcoded human TF open reading frame (ORF) library that includes 3,548 human TF isoforms. TF in this study broadly encompasses sequence-specific activators, repressors, GTFs, chromatin remodelers, etc. Each TF ORF and cell is uniquely barcoded, enabling simultaneous assessment of the transcriptional consequences of overexpressing thousands of TFs in single cells. We hypothesized that systematic shifts in *K_search_* and *K_proof_* would reflect distinct regulatory functions of individual TFs.

To validate the KP model approach, we carried out a large-scale analysis on *K_search_* and *K_proof_*, using filtered TFs from the TF Atlas dataset, comprising 37,528 genes across 1,145,823 single cells [33]. There were 1,836 unique TFs from 3,548 human TF isoforms. Each TF overexpression condition had a different representation; for instance, 49,539 cells were annotated as overexpressing GFP (control), while only 227 cells overexpressed *TBP*. GFP- and mCherry-overexpressing cells served as internal negative controls across all comparisons. Following preprocessing and filtering (see Methods), we retained a subset of genes with matched decay rate information and constructed TF-specific count matrices for downstream kinetic inference. These datasets, together with TBP dwell-time constraints, were used as inputs to the StochasticGene.jl pipeline to infer transcriptional kinetic parameters for each TF overexpression condition (Figure 5A). The resulting kinetic rate estimates were then used to compute *K_search_* and *K_proof_* values per gene and condition.

**Figure 5.**
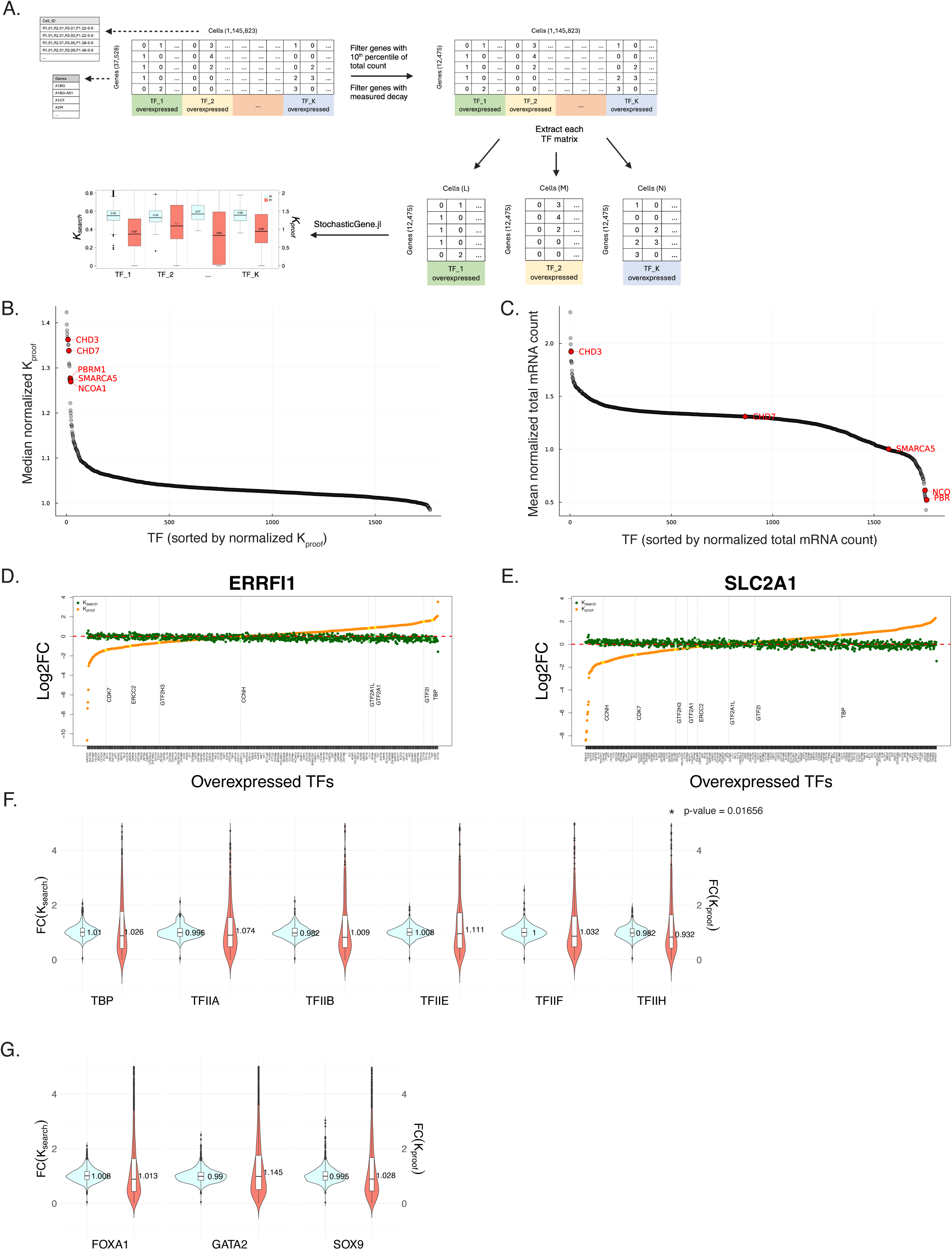
Analysis of TF Atlas using kinetic proofreading model. A) Workflow for simulating TF Atlas dataset using *StochasticGene.jl*. TF-specific effects shown as B) TF vs. Median normalized *K_proof_* and C) TF vs. Normalized total mRNA count, highlighting enrichment of nucleosome and chromatin-remodeling factors (*CHD3*, *CHD7*, *PBRM1*, *SMARCA5*, and *NCOA1*). Gene-specific analyses for D) *ERRFI1* and E) *SLC2A1*, showing log2 fold changes in response to TF overexpression. Subunits of *TFIIH* (*CDK7*, *ERCC2*, *GTF2H3*, and *CCNH*) are highlighted alongside other GTFs. Violin plots summarizing fold change in *K_search_* and *K_proof_* for F) GTFs and G) pioneer factors.

**Figure 6.**
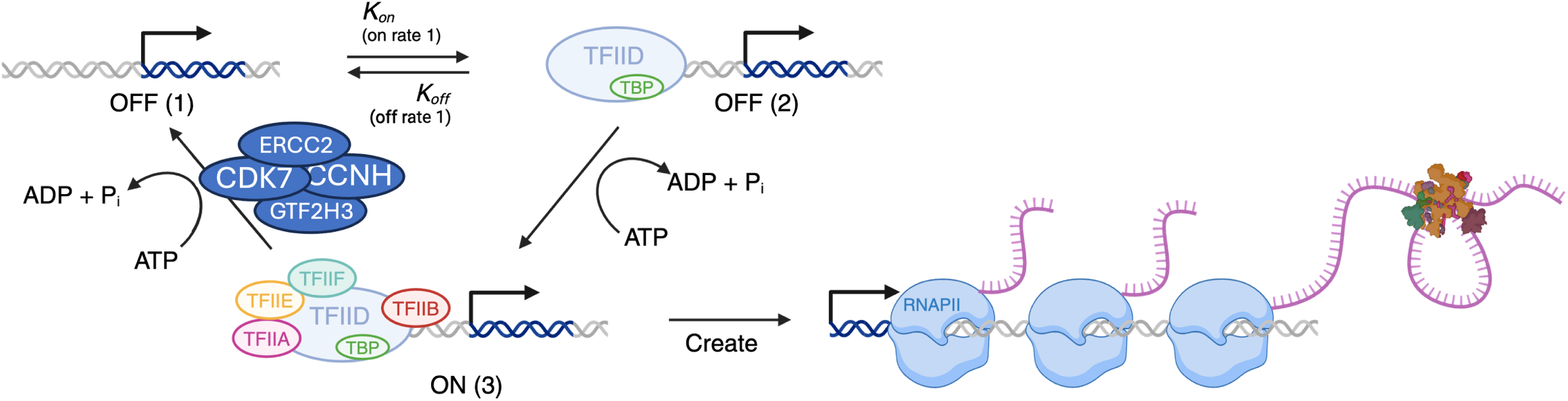
Illustration of kinetic proofreading model. Illustration of the KP model with GTFs, subunits of *TFIIH*, and RNAPII.

This comprehensive analysis provided a global validation of the KP model and revealed several biologically coherent patterns. In particular, when ranking TFs by their global effect on *K_proof_* relative to GFP control, we identified a striking enrichment for chromatin remodelers among the top hits (Figure 5B). The strongest effect was observed for *CHD3*, a core component of the nucleosome remodeling and deacetylase (NuRD) chromatin remodeling complex [91] along with other ATP-dependent chromatin remodelers such as *CHD7*, *SMARCA5* [92–94], *PBRM1* [95], and *NCOA1* [96, 97]. These findings are consistent with the model’s sensitivity to factors that modulate promoter accessibility and nucleosome dynamics. Notably, when we repeated the ranked analysis using simple total RNA fold change over GFP rather than *K_proof_*, the chromatin remodelers are variably distributed and no longer appeared as top-ranked hits, suggesting a consistent kinetic effect that was masked by total RNA level, which is of course the cumulative result of many rates (Figure 5C). This comparison indicates the value added through our modeling of scRNA-seq distributions with the KP model: we identified biochemical activity that would have been missed by just considering total RNA levels.

After validating the approach globally, we next focused on gene-specific analyses to understand why some genes display distinct kinetic features. In particular, we examined *ERRFI1* and *SLC2A1*—two genes previously modeled as 3-state genes in our training set—across 1,836 TF over-expression conditions. This approach inverted the previous analysis: instead of asking how a TF affects all genes, we assessed how the kinetic profile of each gene responds to diverse TF perturbations. For *ERRFI1*, *K_search_* remained relatively stable across all conditions, but *K_proof_* varied markedly. We noticed that GTFs such as *TBP* and *TFIIA* reduced proofreading efficiency (higher *K_proof_*), whereas over-expression of multiple *TFIIH* subunits (*CDK7*, *ERCC2*, *GTF2H3*, and *CCNH*) substantially enhanced proofreading efficiency (lower *K_proof_*) (Figure 5D). Likewise, *SLC2A1* showed a similar response, with lower *K_proof_* under *TFIIH* overexpression (enhanced proofreading efficiency) (Figure 5E). The concordant *K_proof_* trends observed for *ERRFI1* and *SLC2A1* are consistent with both genes being regulated by transcriptional programs that rely on efficient early elongation [98, 99].

These observations illustrate that kinetic proofreading control may be highly gene-specific and dependent on both promoter architecture and co-factor context. Under *TBP* over-expression, for example, *ERRFI1* showed reduced *K_proof_* despite slightly decreased total RNA levels, emphasizing that kinetic modeling captures mechanistic steps not evident from bulk expression alone. Together, these findings suggest that while GTFs such as TBP broadly promote transcription initiation, factors such as *TFIIH*—particularly energy-consuming enzymes such as the CDK7 kinase and ERCC2/XPD helicase subunits—modulate proofreading efficiency in a promoter- and gene-specific manner.

These data also indicated nuanced effects of the GTFs which comprise the pre-initiation complex. To refine and contextualize these observations, we analyzed a focused panel of GTFs (*TBP*, *TFIIA*, *TFIIB*, *TFIIE*, *TFIIF*, and *TFIIH*) and compared to pioneer factors (*GATA2*, *FOXA1*, *SOX9*) to evaluate their impact on transcriptional initiation and proofreading dynamics (Figures 5F and G, respectively). While most GTFs and pioneer factors did not substantially alter *K_search_ or K_proof_* when averaged over all genes, *TFIIH* stood out as consistently decreasing *K_proof_* across many genes, further supporting a role as an ATP-dependent regulator of transcriptional proofreading. Decreased *K_proof_* observed under *TFIIH* over-expression compared with GFP may reflect enhanced proofreading efficiency or a slower transition into the transcriptionally competent state (ON rate 2), suggesting a potential role of TFIIH on the transcription proofreading mechanism. This finding is particularly interesting given that TFIIH has multiple ATP-dependent domains, plays a role in transcription activation by opening the transcription bubble through DNA helicase activity, and is involved in the later stages of transcription initiation [36, 100]. In summary, the hypothesis emerging from the large-scale TF Atlas screen is that chromatin remodelers and components of the general transcription machinery—especially TFIIH—represent key determinants of transcriptional proofreading specificity.

## Discussion

In this paper, we have attempted to integrate multimodal measurements of gene expression all performed in the same cell type under standardized conditions to arrive at a highly constrained kinetic model of single-cell expression. These measurements are: smFISH, scRNA-seq, live-cell RNA imaging, and SMT of GTF TBP. A primary conclusion is that 3-state models show a quantitative improvement over 2-state models, with some genes such as *ERRFI1* showing unambiguous multi-state behavior. We show how these multi-state models, which demonstrate proofreading behavior, have the potential to solve a long-standing question about transcriptional specificity. Finally, we use this highly constrained model to analyze perturb-seq data in an unrelated cell type to infer biochemical mechanisms. Surprisingly, we identify the PIC as a potential player in the proofreading pathway. These predictions are tested in part in an accompanying paper from Kim et al.

### Implications of 3-state models

A unique aspect of our analysis is using SMT of TBP as an input into the model. TBP is a GTF involved in early stages of PIC formation, and the dynamics of TBP have been shown to change with changes in transcriptional bursting [38]. These dynamics are what push the preference for 3- over 2-state models. TBP may be unique in this regard because many genes require TBP based on the structure of their core promoters and thus a tracking measurement at random sites in the nucleus is likely to capture this behavior. In contrast, SMT of upstream transcriptional activators may be dominated primarily by non-specific binding. These observations get to the heart of major difficulty in the field: it is currently impossible to measure TF dynamics at a single regulatory element in the nucleus due to the density of chromatin and the optical resolution of the microscope. Here, computational models can play an essential role by allowing us to infer what these dynamics might look like based on the overall output of gene expression.

The existence of an intermediate state allows the model to have proofreading properties, which we explore theoretically but not experimentally. For a 2-state model to recapitulate some of these properties would require both the ON rate and OFF rate to depend on activator identity. Whether activator search times can show specificity, i.e. the specific activator has a faster ON rate for binding to the cognate motif than the ON rate for a non-specific activator, has not been rigorously tested experimentally. However, we favor an OFF rate-centric view of gene regulation based on a variety of observations in cells. First, gene activity of the GAL genes in yeast was shown to be linearly proportional to the *in vitro* dwell times [37]. Second, for synthetic TFs in human cells where both the ON rate and OFF rate could be independently tuned, only the OFF rate correlated strongly with activity [83]. In our data, the off rate from the intermediate state is much slower than the off rate for initial binding (Tables 1 and 2, comparing *k_p_* and *k_off_*), suggesting that the TF is perhaps stabilized in the context of a PIC. These observations are also consistent with decades of *in vitro* biochemistry experiments.

Interestingly, branched models are not necessary to show proofreading, counter to our expectation. This observation is based on a measure of transcriptional specificity defined herein which we think could be experimentally tractable in live-cell single-molecule imaging. Specificity is defined as the fraction of transcripts produced from binding of a specific activator out of the total number of transcripts potentially produced from the gene. Using this definition, our linear 3-state model and branched KP model both have specificity which can theoretically approach unity. We note that in Hopfield’s original formulation it was simply assumed that the transitions between OFF (2) and ON (3) would be ‘totally insensitive to the difference’ between specific and non-specific pathways, which is an assumption we have relaxed in our treatment. Moreover, by allowing the reverse rates in the 3-state model to both be activator dependent, we are effectively increasing the Gibbs free energy of discrimination and circumventing the need for a branched proofreading mechanism. We find in fact that the linear 3-state model can outperform the branched KP model in certain rate regimes for specificity, however we have opted to use the branched model for the bulk of our analysis to draw explicit parallels to previous work.

Experimentally testing models of transcriptional proofreading is a pressing challenge [68]. Although addition of other sequencing methodologies will certainly constrain models [25, 26], we strongly favor approaches which rely on live-cell imaging, despite limitations in throughput. Previously proposed tests for kinetic proofreading invoke the input response such as precise measures of gene activity with response to concentration of activator [101]. Such validation might be possible by live-cell imaging of nascent RNA accompanied by acute degron depletion of specific TFs. The same study from Tkačik and co-workers also describes how high noise is a consequence of the long tail of extended ON events interspersed with short OFF events which arise from proofreading mechanisms, the combination of which can distort population statistics. Interestingly for *ERRFI1*, the model predicts some ON states due to TF binding which last > 10 minutes (see Table 1), but such events may be < 1% of binding events. Ultimately, the competition framework proposed here further complicates live-cell measurements due to combinations of specific and non-specific binding. The resulting kinetic predictions will be difficult to disambiguate given the dynamic range of cellular imaging and the well-known limitations on resolving multi-exponential distributions [102]. Regardless, there is an emerging consensus that non-equilibrium schemes which describe bursty gene expression and variability are intimately associated with specificity through models of kinetic proofreading [103, 104].

### Inferring biochemical mechanisms from scRNA-seq data

Although scRNA-seq coupled with quantitative modeling has been used to infer kinetic parameters of transcription in the past [87, 105], the rapid growth of scRNA-seq for screening presents a unique opportunity for inferring kinetic biochemical mechanisms. We argue that having a highly constrained model of expression variability will enable better utilization of this data. The basis of our approach is the use of a ground truth data set of 10 genes which we have been characterized in depth to establish strong priors for the steady state measurements. While steady state measurements can never fully substitute time-resolved measurements in cells, we were able to identify two dimension less shape parameters *K_search_* and *K_proof_*, whose values can be recovered from scRNA-seq in our model. These constants reflect TF search efficiency and commitment to productive transcription, respectively. Importantly, *K_proof_* emerged as a sensitive readout of regulatory complexity.

We applied this approach to the TF Atlas dataset and identified potential molecular correlates of our mathematical model. We used the TF atlas analysis both to identify gene-specific regulators of transcription (for example for the *ERRFI1* gene) or the global effects of certain trans-acting factors (for example components of the PIC). As a demonstration of the success of this approach, chromatin remodelers emerged as having strong effects on *K_proof_,* an effect which would have been masked by simply looking at fold change in RNA. One interesting hit was *TFIIH*. TFIIH is a multifunctional complex involved in promoter opening via DNA helicase activity [36, 96]. When correctly assembled and positioned at the promoter by a sequence-specific factor, TFIIH facilitates productive transcription initiation by unwinding the DNA duplex to form the transcription bubble. However, TFIIH has also been implicated in the turnover of TFs and GTFs binding to chromatin. For example, the CDK7 kinase component of TFIIH directly phosphorylates androgen receptor, leading to proteolytic turnover of this ligand-regulated TF, which is essential for transactivation of target genes [106]. Our observations directly support this mechanism. However, we further speculate if TFIIH is mispositioned or recruited by a nonspecific factor, its helicase/kinase activity may still function—but in a destructive manner. Instead of promoting initiation, it may destabilize the PIC and displace GTFs, effectively returning the gene to the first OFF state. Our modeling results support this possibility: overexpression of multiple TFIIH subunits led to decreased *K_proof_* in certain genes, suggesting that excess TFIIH may increase proofreading activity rather than simply promoting transcription. This prediction, that overexpression of GTFs like *TFIIH* can enhance transcriptional proofreading, is experimentally testable and highlights a potential mechanism of gene-specific regulation through kinetic gating.

In summary, we present a robust framework for inferring transcriptional kinetics by integrating orthogonal single-cell data modalities. The KP model allows for a mechanistic interpretation of TF function in terms of both search and proofreading dynamics and provides a scalable strategy for functional analysis in genomic and perturbation datasets. Future work should aim to validate these predictions using targeted live-cell measurements and chromatin-level assays, and to expand this framework to other TFs, cell types, and regulatory contexts.

## Acknowledgements

We thank members of Larson laboratory at NCI for helpful discussions and insightful comments. This work utilized the computational resources of the NIH HPC Biowulf cluster (https://hpc.nih.gov). PT was supported by the National Institute of Arthritis and Musculoskeletal and Skin Diseases Intramural Research Program. The contributions of the NIH author(s) were made as part of their official duties as NIH federal employees, are in compliance with agency policy requirements, and are considered Works of the United States Government. However, the findings and conclusions presented in this paper are those of the author(s) and do not necessarily reflect the views of the NIH or the U.S. Department of Health and Human Services.

## Figure Legends

Figure S1. Posterior distributions of the fitted rates

Posterior distributions of the fitted rates for the 10 genes: *CANX, DNAJC5, ERRFI1, KPNB1, MYH9, RAB7A, RHOA, RPAP3, SEC16A,* and *SLC2A1*. Red dashed line represents the maximum likelihood of the fitted rates, and blue dotted line indicates the median of the fitted rates.

Figure S2. Comparison between specific and non-specific TF binding in the KP model

Illustration of the KP model with non-specific (ns) and specific (s) binding of TFs.

## Methods

### Rate inference using StochasticGene.jl

We fit 2-state, 3-state, and KP models to the multimodal dataset for each gene using a previously published stochastic model [19, 21]. All model fitting and simulations were performed with StochasticGene.jl (freely available at https://github.com/nih-niddk-mbs/StochasticGene.jl/) version 1.1.7 and the Julia language version 1.10.4. To minimize the number of parameters, which allowed for faster convergence and greater identifiability, we fixed R = S = 3 with all elongation rates set to the same value r and all splicing rates set to the same value s (where r ≠ s). We found that increasing R beyond 3 rarely improved model performance given the uncertainty in our data (data not shown). In order to constrain the model to fit both kinetic and steady state data, we used TBP as the universal TF and constrained the priors for splicing event to 0.05 per minute. The convergence of the simulation was assessed by three criteria: (1) maintenance of the MCMC acceptance rate (fraction of proposed parameters accepted by the sampler) to within the optimal range of 20-40% [107, 108]; (2) ensurance that the maximum likelihood estimates converged to a steady value; and (3) iterating for long enough such that *r̂* (rhat) was less than 1.1 [109]. After each iteration, the initial and final maximum log-likelihood values were compared; if they differed, the sampling continued until the convergence criteria were met.

### Competition Model

Define a transcriptional network where a single gene is present in the milieu of the nucleus where many different transcription factors (TFs) are present. For this model, the gene is activated by binding of a specific factor S but other non-specific (NS) transcription factors may also bind and/or activate the gene. It is expected these NS factors will activate the gene with lower efficiency, but this assumption is not necessary. The branched kinetic proofreading scheme is displayed, but any kinetic rate model is applicable. In general, transcriptional activation proceeds through a series of steps which may or may not occur with similar rates for S and NS. The general assumption is that transcription is initiated by binding of some sort of activator with sequence preference (“single target factor”), followed by recruitment of general transcription factors (GTFs) which work across many genes (“multitarget factors”), and finally initiation of transcription by RNA polymerase II. The model is agnostic to the molecular identity of these states and says nothing, for example, about whether binding sites are in the enhancer or the promoter, or whether pre-initiation complex recruitment or pause release or some other step is rate-limiting.

#### State Description

*A*: a single gene in the unoccupied state with neither *S* nor *NS* bound.

*B_S,NS_*: the same gene now occupied either by a *S* or *NS*. This occupation would correspond to binding of an activator with sequence preference which is capable of initiating downstream steps such as GTF recruitment.

*C_S,NS_*: an intermediate state which occurs after initial binding of the activator. This state could correspond to GTF recruitment (mediator, TBP, etc.), enhancer looping, or any manner of intermediate state. Conceptually, there is likely overlap between the molecules recruited to constitute the *C_S_* and *C_NS_* states, although this commonality is not explicit in the model.

*R_S,NS_*: the state which is committed to RNA synthesis. Whether this state is considered the post-pause release nascent RNA polymerase II or completed mRNA will depend on the details of the experimental system. For simplicity in description, we will refer to *R* as ‘RNA’ or ‘transcripts’, meaning nascent RNA transcripts which are fully synthesized into mRNA.

This system is capture by the following system of inhomogeneous differential equations, where *Init* is the flux into the system:

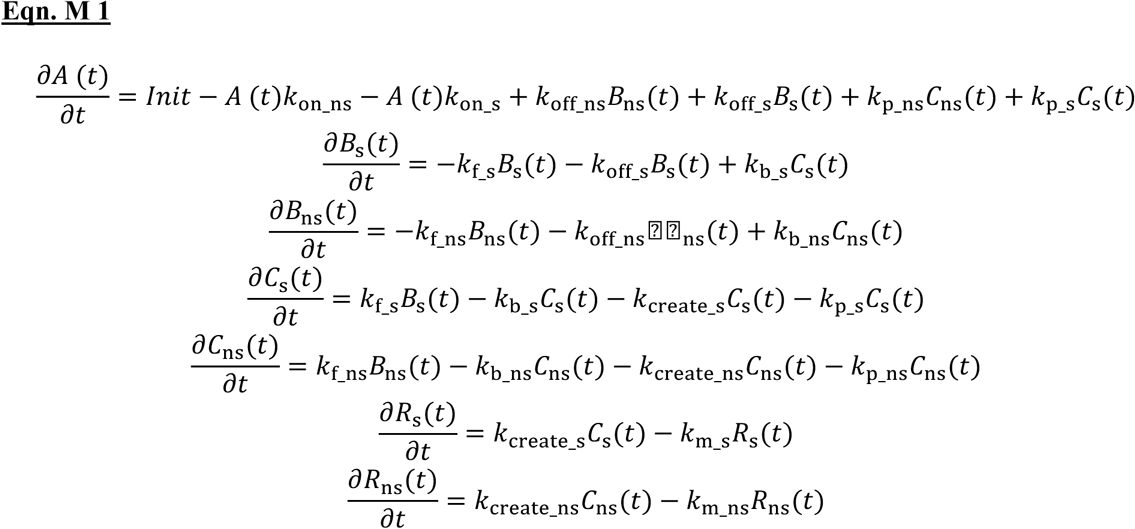

First, we treat the steady state case, where the left side of each equation is 0. This system can be solved using standard linear algebra techniques.

#### Derivation of the specificity formulas

We define the specificity as:

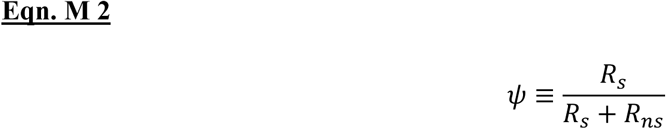

Which is the ratio of all transcripts produced through the specific pathway (the *R_s_* branch of the pathway, depicted on the right side) to all transcripts produced through both pathways (*R_s_+R_ns_*). Biologically, this quantity would correspond to the fraction of all transcripts resulting from binding of a particular TF out of the total number of transcripts produced from the gene. The particular form of the equation is chosen to resemble the commonly used formula for analyzing RNA splicing isoform distributions, which is “percent spliced in” p.s.i or PSI (ψ Indeed, there are strong similarities between the mathematical results for specificity in transcription and specificity in splicing.

The full formula for ψ is given here:

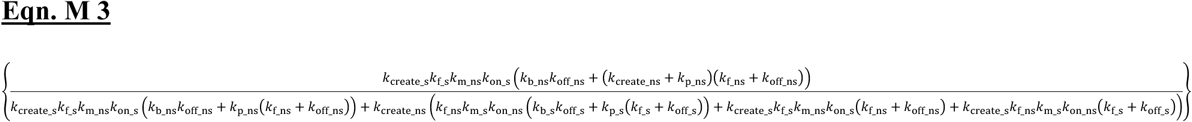

#### Limiting cases

We now treat several limiting cases of this formula.

#### Symmetrical rate constants

If *k_off_s_ = k_off_ns_, k_on_s_=k_on_ns_, k_create_s_=k_create_ns_, k_b_s_=k_b_ns_, k_p_s_=k_p_ns_,* each outcome is equally likely. Basically, there is no order of encounter, and both transcripts are at equal abundance. This simple check gives ψ = ½.

#### Telegraph model

In the telegraph model, the only off rates are *k_off_s_* and *k_off_ns_*. Thus*, k_b_s_=k_b_ns_=k_p_s_=k_p_ns_*=0. *k_f_s_* and *k_f_ns_* play the role of irreversible forward rates, and the downstream forward rates do not appear in the final result. The result is:

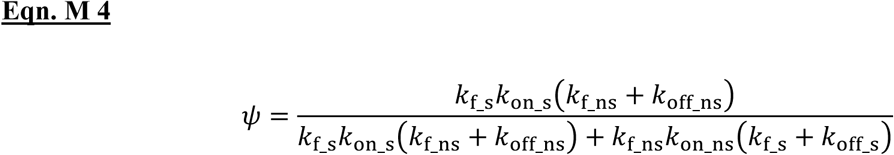

Which is also Eqn. 2 above.

We now set *k_f_s_=k_f_ns_=k_f_* which is equivalent to saying that the irreversible forward rate does not vary between specific and non-specific pathways. Biochemically, this assumption is equivalent to saying that once the GTFs are recruited (say in assembly of a pre-initiation complex) the forward rate to initiate RNA synthesis is the same regardless of which activator was initially bound. The reasoning behind this assumption is that TFs are all capable of initiating transcription at certain genes, with the specificity coming from binding cognate DNA binding sites. The result is then:

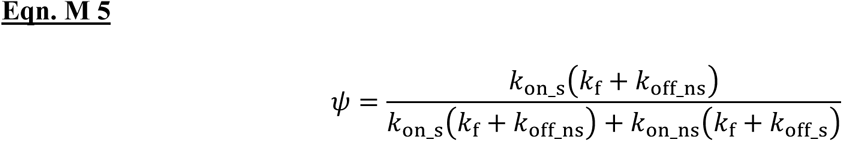

Consider now the case of differing off rate between specific and non-specific TF but identical on rates. This case would correspond, for example, to two TFs with identical expression levels and on rates -- they both have equal likelihood of encountering the binding site – but the specific factor has (for example) a longer dwell time on the DNA binding site. The specificity then reduces to:

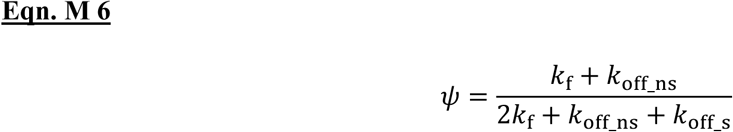

In the limit where *k_f_* is large compared to *k_off_s_*, corresponding to very fast forward progression:

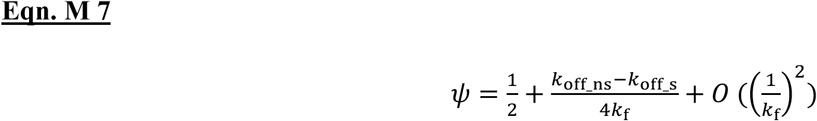

Which is Eqn. 3 above. The specificity is ½ to first order: because *k_f_* is fast, most binding events lead to transcription and are therefore locked in. The specificity improves with the difference between specific and non-specific binding with respect to this otherwise large *k_f_*.

However, this limiting case does not capture the actual biological situation correctly. In the nucleus, there are likely many more non-specific TF than specific TF. Thus [NS] >>[ S]. Going forward, we capture this biological reality by making *k_on_ns_* >> *k_on_s_*. For the sake of simplicity, we assume that there are M-fold more non-specific factors but they bind M-fold less tightly. Also, we draw on the observation that transcription is quite inefficient: many TF binding encounters do not lead to transcription initiation. Thus, the limits are: 1) *k_f_* is small compared to *k_off_s_*, corresponding to many unproductive binding events, and 2) M >>1, corresponding to the specific factor binding more tightly than the non-specific factor but being much less abundant:

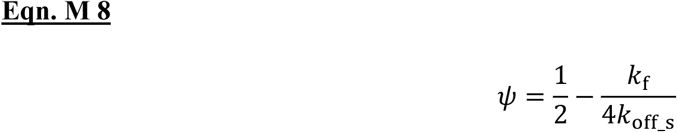

Which is Eqn. 4 above. The primary conclusion is that the maximum specificity provided by the telegraph model is ½. Intuitively, in the case where non-specific factors are M-fold more abundant but bind with M-fold shorter dwell time, the effective occupancy of the regulatory element by non-specific TFs is the same as the occupancy by the specific TF, making distinguishing these two scenarios impossible: if specificity is only encoded in the off rate, the gene does not distinguish specific from non-specific binding.

#### Branched Proofreading Model

We continue this analysis now for the full branched proofreading model depicted in Figure S2. We add the critical feature of a proofreading model, which is that the proofreading off rate (*k_p_*) is also sensitive to the specific and non-specific pathways. In other words, the activator identity is sensed twice. To investigate the effects of the specific topology of the model (see also below), we set *k_bs_ = k_b_ns_ =0*. So, the two off rates in the system are only through initial binding (*B* state) and disassembly of the intermediate state (*C* state). Again, for simplicity, we assume that the *k_p_* is M-fold slower for the specific factor than the nonspecific factors. In this case, the specificity is:

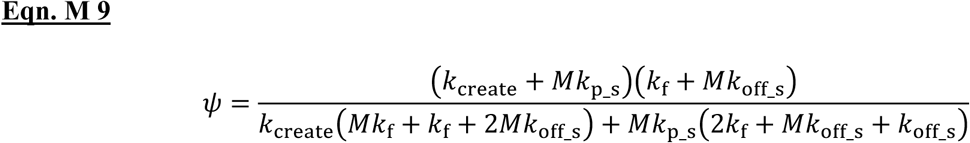

We also continue with the same assumptions which likely approximate the biological situation: 1) *k_f_* and *k_create_* are small compared to *k_off_s_*, corresponding to relatively weak forward rates and many unproductive binding events, and 2) M >>1, corresponding to the specific factor binding more tightly (now at two different steps in the kinetic model) than the non-specific factor but being much less abundant.

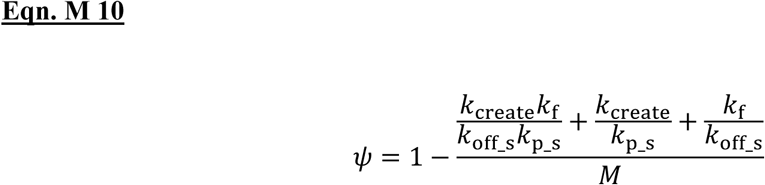

The primary result is that the maximum accuracy is now 1. In the proofreading scheme, which introduces two TF-dependent off rates, arbitrarily high specificity levels can be achieved. The above result is particularly suited to transcriptional activation, where there are many unproductive events.

#### Linear Proofreading

The linear model has the same number of rates (forward and backward) as the branched proofreading model, and the only difference is the topology. The branched proofreading model has a C → A transition, and the linear proofreading model has C→B transition. The former is consistent with the traditional proofreading kinetic scheme and indeed is usually asserted to be necessary. Biochemically, this C→ A step would correspond to an irreversible, energy-dependent step where DNA-bound complexes are completely removed, thereby resetting the gene. While in the linear model, this off rate from the intermediate state is back to the B state, i.e. the activator bound state. The same assumptions are applied for the branched kinetic proofreading model.

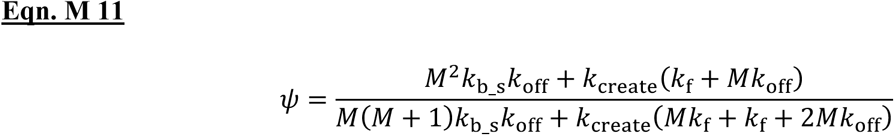

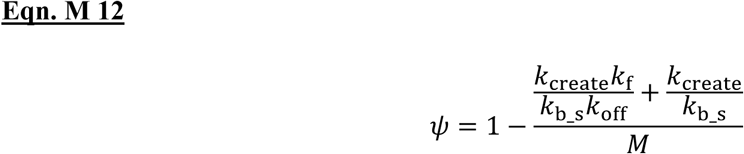

For the linear model, the maximum specificity is again 1, meaning that arbitrarily high accuracy is possible with 2 activator-dependent off rates.

#### Comparing accuracy of the linear and branched models

Interestingly, when comparing Eqns. 10 and 12, it appears that the linear model returns higher specificity (*ψ*) because there are fewer negative terms. To check when this observation is true, we compute the difference Δ*ψ* = *ψ_branched_* – *ψ_linear_*. We set all *k_b_* and *k_p_* rates to a generic *k_off_int_*, meaning the off rate from the intermediate complex. In this manner, we interrogate the role of the kinetic structure rather than the particular value of the off rate. We set the on rate of the non-specific pathway to be P-fold greater than the specific pathway. The result for Δψ is:

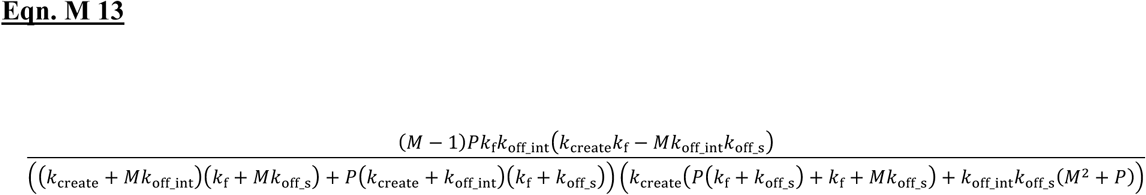

Since the denominator is positive, and M>1 by definition, there is a clear change of sign at:

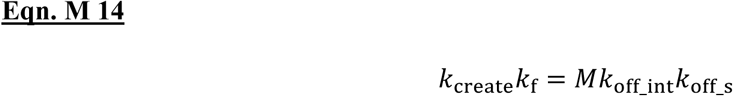

When the product of forward rates is greater than the product of backward rates, ψ_branched_ > ψ_linear_, and ψ_branched_ will provide greater specificity. This result is consistent with the original proofreading models which maintained that the branched pathway was needed to avoid the buildup of abundant intermediates which would drive the reaction forward by mass action, even for slow kinetic steps. The branch transition (C→A) required the reaction to go back to the unoccupied state to circumvent this build up of an intermediate state. In contrast, if the product of the backward rates is greater than the product of the forward rates, then ψ_branched_ < ψ_linear_, and ψ_linear_ will provide greater specificity. We speculate this regime is the more relevant for transcription, since many TF binding events are unproductive. In this case, the branched pathway is not necessary because the intermediate state quickly returns to the unoccupied state anyway (C → B →A).

In summary, both the classic branched proofreading model and a linear proofreading model provide a theoretical specificity of ψ=1. It may be there are biochemical reasons to prefer one over the other, but mathematically, the linear model provides greater specificity under the rate conditions likely to prevail for transcriptional activation.

### scRNA-seq processing and quality control

scRNA-seq data from HBECs were generated using the 10x Genomics platform and processed with the Cell Ranger pipeline (version 3.0.0). Sequencing produced a total of 4,074 cells with a mean of over 98,000 reads per cell and a median of over 4,100 detected genes per cell. Of all reads, 79.7% mapped to the genome and 64.9% were confidently assigned to a unique gene in hg19 transcriptome. Initial processing yielded a count matrix comprising 32,728 genes across 4,074 cells. Quality control filtering was applied to remove low-quality cells based on the following criteria: (1) cells with 3,500-5,000 detected genes, (2) cells with 15,000-32,000 detected molecules (UMIs), and (3) mitochondrial transcript fraction below 5%. After filtering, 14,244 genes across 2,879 high-quality cells were retained for downstream analysis (Figure 4A).

### Data collection

The scRNA-seq data used in this study for TF Atlas were obtained from previous studies [33]. Raw count matrix was available and downloaded from the Gene Expression Omnibus database (http://www.ncbi.nlm.nih.gov/geo/; GSE217460). Since the required input for the StochasticGene.jl package is raw counts of a gene per cell, no normalization was required.

### Processing TF Atlas scRNA-seq data and kinetic inference pipeline

scRNA-seq data from the TF Atlas dataset [33] were processed to enable large-scale kinetic modeling. From the full count matrix, genes with zero expression across all cells were removed, along with those falling below the 10th percentile of total gene counts. Genes lacking experimentally measured decay rates were also excluded. This filtering resulted in a refined dataset of 12,475 genes across 1,145,823 cells. To incorporate gene-specific degradation rates into kinetic modeling, we matched this gene list with transcript half-life data previously inferred in HCT116 cells [87]. For each TF of interest, we generated a TF-specific subset of the total count matrix by selecting cells annotated with the corresponding overexpression barcode (Figure 5A). Of the 1,836 unique TFs present in the dataset, 1,773 TF-specific matrices passed filtering and were used for downstream analysis. Each TF-specific dataset, together with matched decay rates and TBP dwell time constraints, was used as input to the StochasticGene.jl pipeline for inference of kinetic parameters under each TF overexpression condition. This large-scale inference required > 17×10^6^ hours of cumulative CPU time on the NIH Biowulf High Performance Cluster.

## Notes

### Competing Interest Statement

The authors have declared no competing interest.

### Summary of Updates

Revised line 442 and 443 where the comments accidentally embedded in the main text.

