## Supplementary figures and images for "Highly Constrained Kinetic Models for Single-Cell Gene Expression Analysis"

### Supplemental Figures

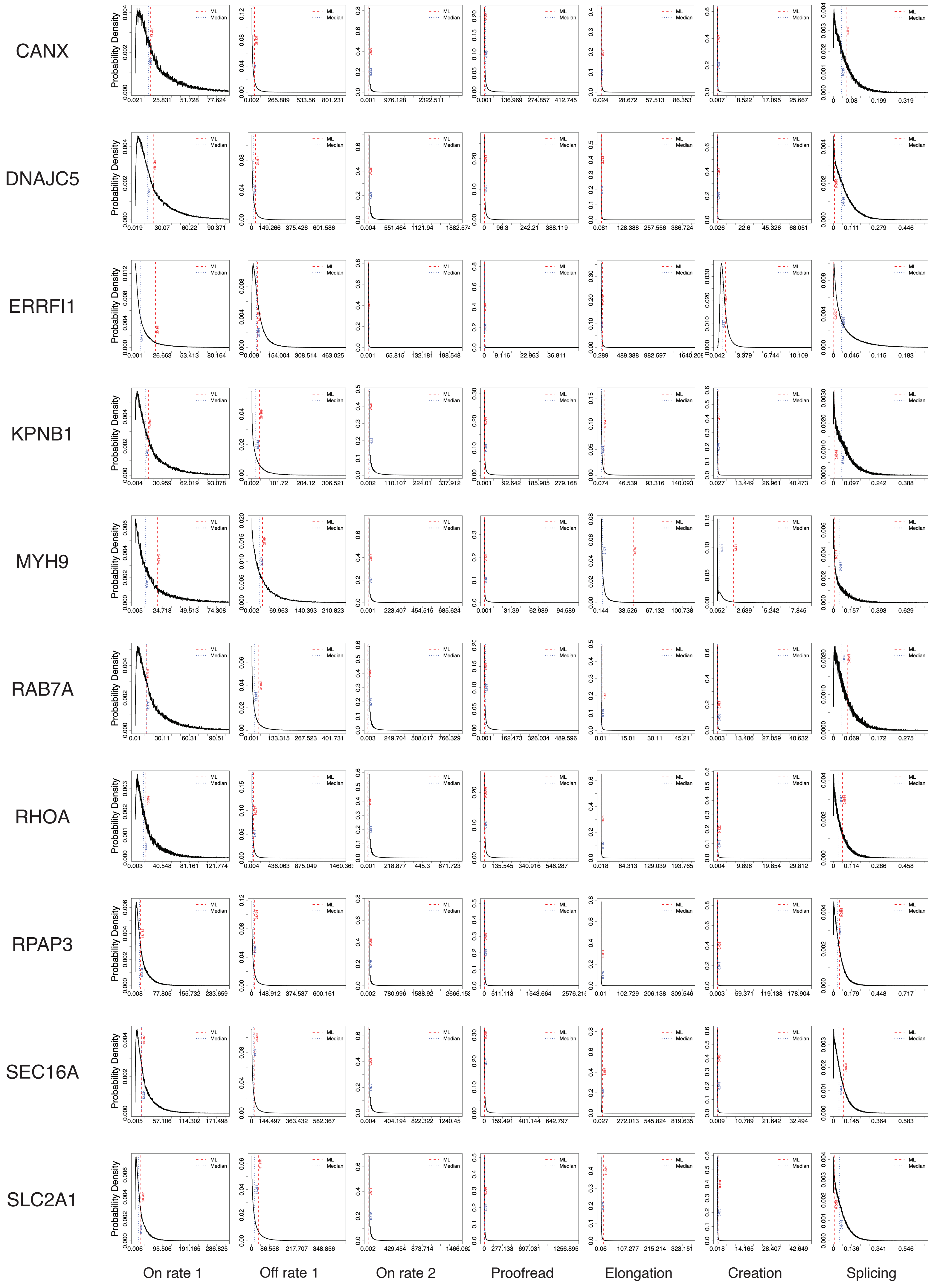

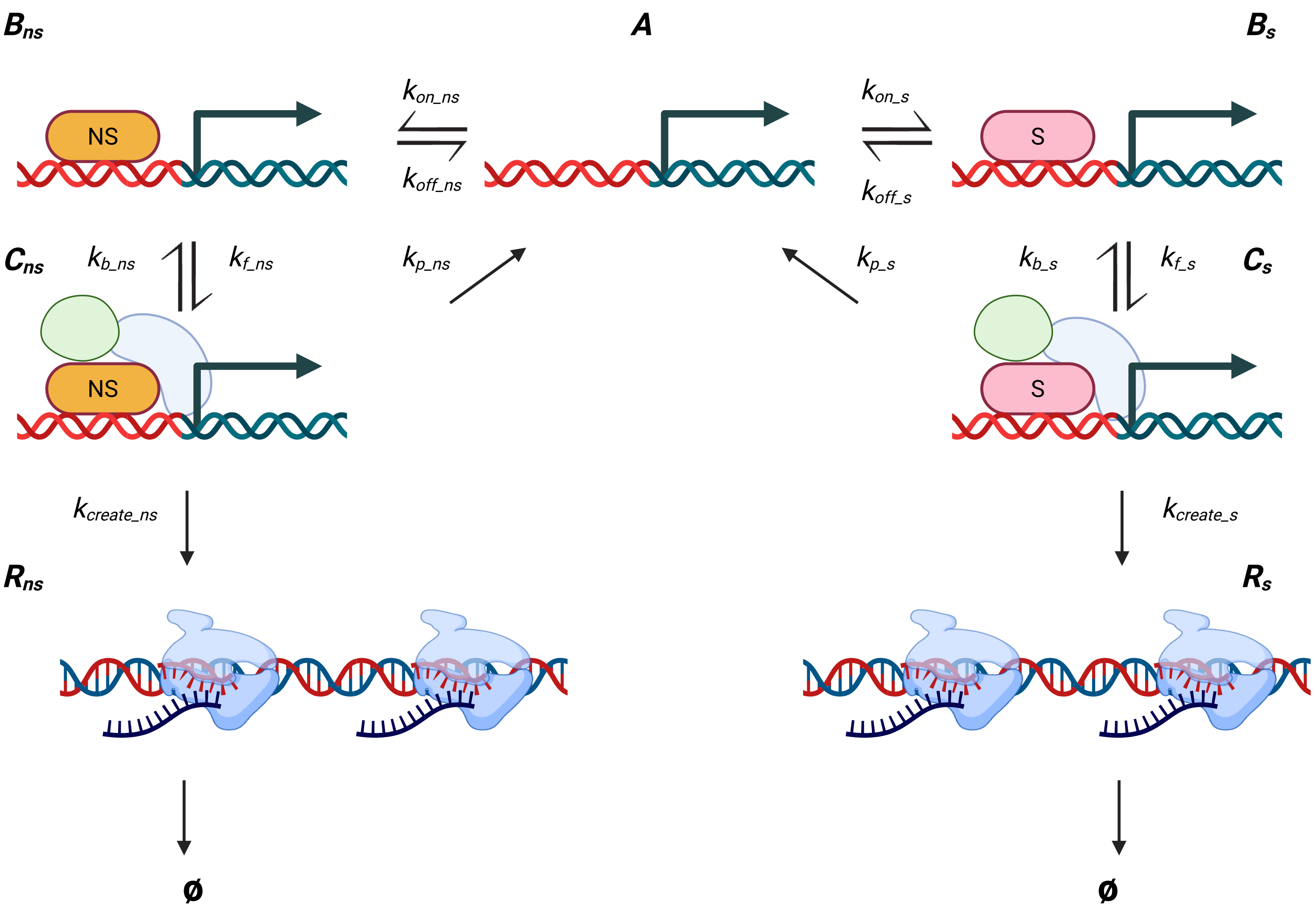
